# Glucocorticoid receptors in oligodendrocyte precursor cells regulate hippocampal network plasticity and stress-induced behavior in mice

**DOI:** 10.1101/2024.12.20.629639

**Authors:** Lorenzo Mattioni, Giulia Poggi, Celine Gallagher, Katrin Becker, Linh Le, Maja Papic, Jasmin Engbers, Maija-Kreetta Koskinen, Ali Abdollahzade, David P. Herzog, Leonardo Nardi, Andrea Conrad, Sarah Winterberg, Christa Merte-Grebe, Liana Melo-Thomas, Hyonseung Lee, Hans Schwarzbach, Jennifer Klüpfel, Ralf Kinscherf, Jan Engelmann, Beat Lutz, Ari Waisman, Iiris Hovatta, Thomas Mittmann, Michael J. Schmeisser, Marianne B Müller, Giulia Treccani

## Abstract

Glucocorticoid receptors (GRs) are key mediators of how the stress hormone glucocorticoids (GCs) shape postnatal brain development and adaptive plasticity. Because GC signaling is critical during this period, postnatal GC concentrations are tightly regulated in the brain, whereas excessive levels of circulating GCs can disrupt developmental trajectories and increase the risk of psychiatric disorders later in life. GR function influences multiple neural cell types, but its cell-specific roles, particularly early in development, remain poorly understood. Oligodendrocyte precursor cells (OPCs), which generate myelinating oligodendrocytes and actively modulate neuronal networks, express GRs and can therefore respond to fluctuations in GC levels. Although excessive GC exposure during early life adversity has been linked to changes in OPC development, the physiological role of GR signaling specifically within OPCs remains unclear. To address this, we conditionally deleted GRs in postnatal OPCs in mice to investigate the role of physiological GC signaling in OPC proliferation and maturation, as well as in neuronal network activity and behavior. This deletion resulted in reduced oligodendrocyte and myelinated axon density in the hippocampus, sex-specific alterations in hippocampal activity and long-term potentiation following acute challenge, and impairments in memory formation in adulthood. Our findings reveal a novel, OPC-specific role for GRs and suggest that physiological GR activity in the oligodendrocyte lineage contributes to normal hippocampal plasticity, learning and memory.

**Significance Statement:** Glucocorticoid receptors (GRs) mediate the effects of the stress hormone glucocorticoids (GCs) on postnatal brain development and adaptive plasticity. While the function of GRs in neurons is well characterized, much less is known about their role in oligodendrocyte precursor cells (OPCs). OPCs, which give rise to myelinating oligodendrocytes and participate in the modulation of neuronal networks, express GRs and can therefore sense fluctuations in GCs during stress response; however, the physiological role of GRs in OPCs remains unclear. In this study, we found that deleting GRs in early postnatal OPCs reduced the density of oligodendrocytes and of myelinated axons, altered hippocampal activity and long-term potentiation in response to acute challenge, and impaired memory formation in adult mice. These findings identify OPCs as key targets of GR signaling and suggest that physiological receptor activity in the oligodendrocyte lineage contributes to normal hippocampal plasticity, learning, and memory.

## Introduction

The early postnatal period is a critical developmental stage, when exposure to hormones influences brain maturation (1). Glucocorticoids (GCs), the main hormones released in response to stress, are tightly regulated during this time of development that is defined as the stress-hyporesponsive period (SHRP) and is characterized by reduced secretion of GCs following mild stress conditions (2). Although exposure to negative stress in early life can override the SHRP and expose the brain to harmful surges in GCs, endogenous GCs are also present postnatally and play a beneficial role in brain development (3). Nevertheless, the specific cellular effects of GCs on postnatal brain maturation remain poorly understood.

Importantly, different cell populations of the brain are endowed with glucocorticoid receptors (GRs), making them sensitive to physiological fluctuations, but also to stress-induced increase of GCs (4). One glial cell population that has been identified both clinically and preclinically as an emerging player in stress-related psychiatric disorders are oligodendrocyte precursor cells (OPCs) (5–9). OPCs express several markers, including the chondroitin sulfate proteoglycan 4 (CSPG4) or neuron-glial antigen 2 (NG2), and are therefore also referred to as NG2-glia (10, 11). They are homogeneously distributed throughout the CNS, both during development and adulthood, and they are highly responsive to external stimuli, including neuronal activity (12). Through their canonical pathway, OPCs can give rise to myelinating oligodendrocytes. However, a large proportion of OPCs remain undifferentiated and survey the surrounding extracellular environment (13). These resident OPCs are thought to play a critical role in modulating and fine-tuning neuronal networks, through non-canonical (and myelination-independent) pathways (14–18) and are indeed integrated into the neuronal network via synapses with neighboring neurons (19–21). Mice exposed to maternal separation (a model of early life adversity (ELA)) have an increased number of mature oligodendrocytes at postnatal day (PD)15 and a reduction in OPCs as adults (22). In post-mortem studies on individuals exposed to ELA, increased numbers of mature myelinating oligodendrocytes (OLs) and decreased numbers of OPCs were found (23). Taken together, these findings suggest enhanced maturation of OPCs as a result of their early exposure to excess GCs. Recently, we have shown that ELA not only alters the density of OPCs or OLs, but has a direct effect on the OPC gene expression patterns and their electrical properties and that these effects are largely GR-dependent (24).

OPCs remain responsive to stress signals throughout the adult lifespan (25–27) and express GRs in both grey and white matter regions (28–30), indicating that GCs may play a physiological role in the regulation of OPC function and differentiation (31). Although there is accumulating evidence for the influence of GCs on OPC proliferation and maturation, the specific contribution of constitutive GR signaling in OPCs to neuronal network function and behavior remains unclear. Therefore, the aim of this study was to determine how physiological developmental GR expression in OPCs regulates their proliferation and differentiation, and whether GRs in OPCs contribute to modulating hippocampal network activity and fine-tuning of behavioral phenotypes, including those behaviors induced by stress in adulthood. We conditionally deleted GRs in OPCs during the first postnatal week, a developmental window characterized by peak OPC proliferation (32), and examined the long-term consequences on oligodendrocyte dynamics, hippocampal network excitability, and adult behavior. Our findings reveal that postnatal GR signaling in OPCs is required for OPC maturation and axon myelination, fine-tuning of neuronal network potentiation in responses to challenge, and the establishment of adult memory.

## Results

### Conditional deletion of GR in OPCs impairs lineage maturation and density of myelinated axons without affecting proliferation and morphology

To investigate the role of an early conditional knockout (cKO) of GR on OPC differentiation, morphology, as well as behavioral phenotypes in adult mice, we generated an OPC-specific GR cKO mouse strain. We obtained the cKO mice by crossing the Ng2CreER^T2^ mice with Nr3c1^fl/fl^ mice and inducing recombination by tamoxifen injection at PDs 2 and 4 (Fig. 1 *A*; Materials and Methods). As previously reported (28, 30), we confirmed by fluorescence-activated cell sorting (FACS) analysis that GRs are expressed by the hippocampal and cortical OPCs and mature oligodendrocytes (OLs), with greater concentration in OPCs (*SI Appendix*, Fig. S1 A, B). We then assessed the efficiency of GR deletion in OPCs by performing immunofluorescence (IF) staining in the dorsal hippocampus (Fig. 1 *B*) and the dorsally located somatosensory cortex of the cKO and control littermates. Immunostaining revealed a decrease in the percentage of GR+ NG2+ Olig2+ OPCs in cKO mice in the hippocampus (Fig. 1 *C* and *D*) and cortex (*SI Appendix,* Fig. 2 B), where there were no sex differences (*SI Appendix,* Fig. S3 A and B). To confirm the specificity of GR deletion in OPCs, we performed co-staining of GR with several cell-specific markers, including Glial fibrillary acidic protein (GFAP) for astrocytes (*SI Appendix*, Fig. S4 A-C), ionized calcium binding adaptor molecule 1 (Iba1) for microglia (*SI Appendix*, Fig. S4 D-F), NeuN for neurons (*SI Appendix*, Fig. S4 G-I) and Platelet-derived growth factor receptor beta (PDGFRB) for pericytes (*SI Appendix*, Fig. S4 J-L). We did not observe a reduction of GRs in any of these cell populations in cKO mice, either in the hippocampus or in the cortex (*SI Appendix*, Fig. S4).

**Figure 1.**
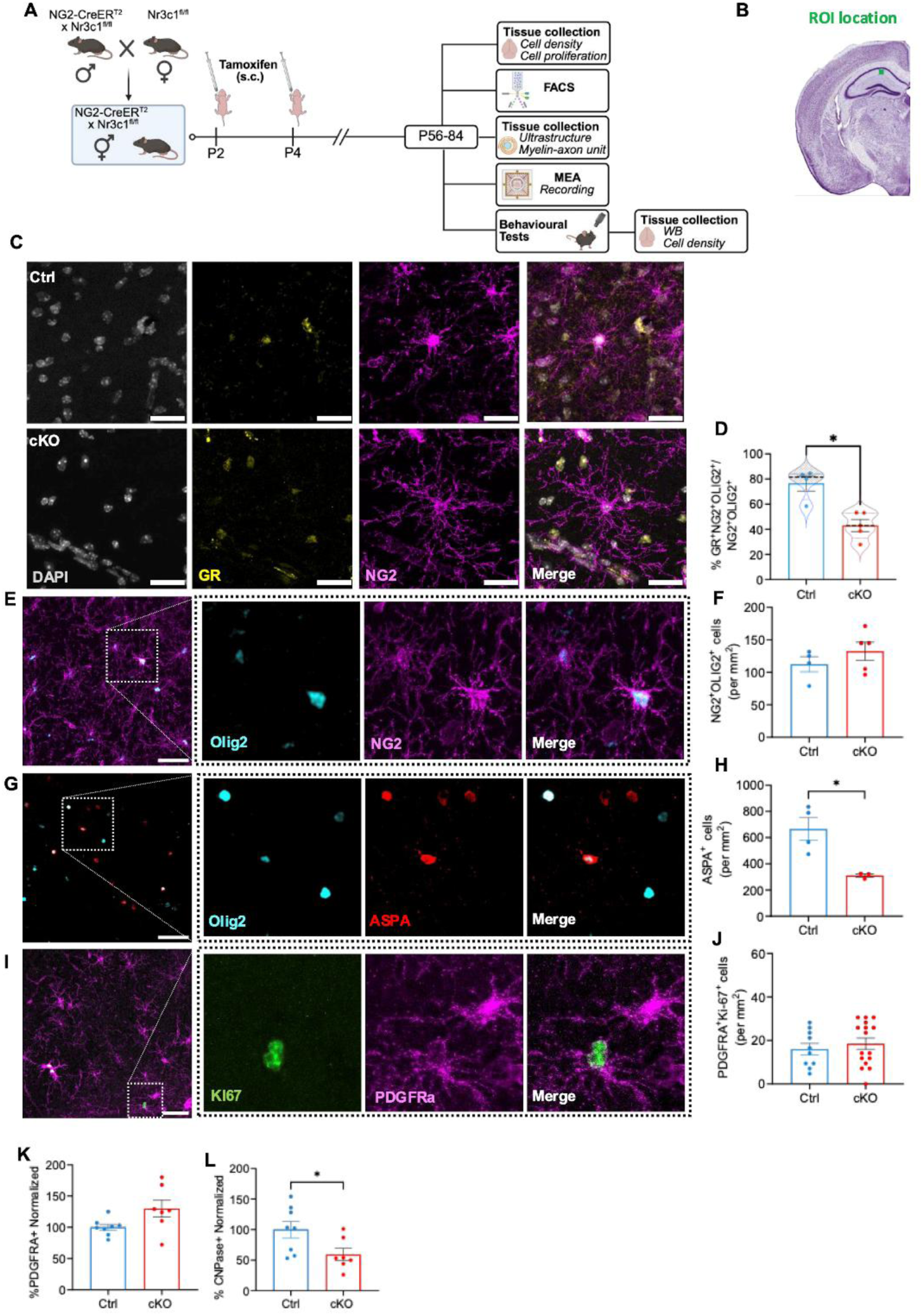
OPCs-specific GR cKO alter oligodendrocyte density but does not induce changes in OPC density and proliferation. (**A**) Breeding strategy and experimental timeline for tamoxifen injections, behavioral studies, MEA recordings, and immunohistochemical analysis. (**B**) Region of interest (ROI, Cornu ammonis 1 (CA1), Bregma: −1.79 to −2.53 mm, image adapted from Paxinos and Franklin mouse brain atlas). (**C-J**) **Immunofluorescence in the hippocampus**. (**C**) Representative confocal images of the co-expression of GR (yellow) and NG2 (magenta) in the hippocampus of Ctrl and cKO mice. Scale bar = 20 µm (**D**) Percentage of GR+NG2+ cells (Mann Whitney test: U = 0, p = 0.0159; n = 4 Ctrl and n = 5 cKO mice). Representative confocal images with digital magnification of the boxed area and related quantification for (**E-F**) NG2 (magenta) and (**G-H**) ASPA (red), each co-stained with Olig2 (cyan). Scale bar = 20 µm. (**F**) Density of NG2+Olig2+ cells (unpaired t test: p = 0.3202; n = 4 Ctrl and n = 5 cKO mice). (**H**) Density of ASPA+ cells (unpaired t test with Welch’s correction: t(3.129) = 4.038, p = 0.0252; n = 4 Ctrl and n = 5 cKO mice). (**I**) Representative confocal images with digital magnification of the boxed area and related quantification for PDGRFA + (magenta) and Ki-67+ (green) cells. Scale bar = 20 µm. (**J**) Density of PDGRFA+Ki67+ cells (unpaired t test: p = 0.5153; n = 10 Ctrl and n = 16 cKO mice). Brightness and contrast of the micrographs have been adjusted for display purposes. (**K-L**) **Flow cytometry in hippocampal tissue.** (**K**) Percentage of PDGRFA+ cells in the CD1145-CD11b- population (unpaired t test with Welch’s correction: p = 0.073; n = 8 Ctrl and n = 7 cKO mice). (**L**) Percentage of CNPase+ cells in the CD1145-CD11b- population (unpaired t test: t(13) = 2.384, p = 0.0331; n = 8 Ctrl and n = 7 cKO mice). Data are expressed as the mean ± S.E.M (with graph D also including an overlaid violin plot for median and data range) and each dot represents one animal. *P < 0.05.

**Figure 2.**
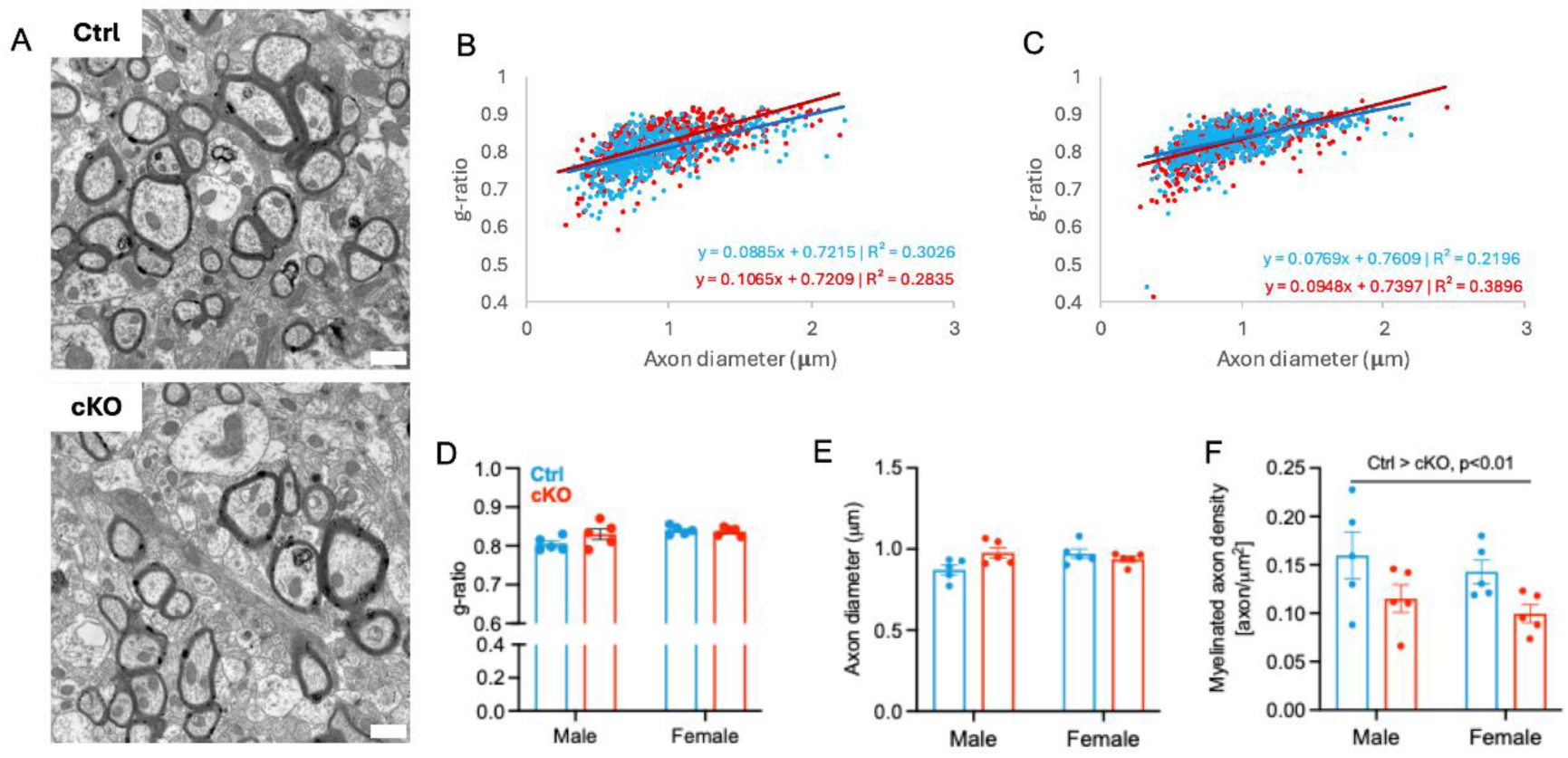
OPC-specific GR cKO reduces the density of myelinated axons in the hippocampus of male and female mice. (**A**) Representative transmission electron microscopy micrograph from Ctrl vs. cKO male, scale bar = 1 µm. (**B**) Relation between g-ratio and axonal diameter in Ctr and cKO male mice. (**C**) Relation between g-ratio and axonal diameter in Ctr and cKO female mice. (**D**) g-ratio (Two-way ANOVA, genotype: F(1, 16) = 1.559, p = 0.2298; sex: F(1, 16) = 5.878, p =0.0275; genotype x sex: F(1, 16) = 2.760, p=0.1161). (**E**) Axon diameter (Two-way ANOVA, genotype: F(1, 16) = 1.520, p=0.2354; sex: F(1, 16) = 1.131, p=0.3034; genotype x sex: F(1, 16) = 6.245, p = 0.0237. Tukey’s multiple comparisons test, n.s.). (**F**) Density of myelinated axons (Two-way ANOVA, genotype: F(1, 16) = 7.468, p=0.0147; sex: F (1, 16) = 1.028, p=0.3258; genotype x sex: F(1, 16) = 0.001840, p=0.9663). Data are expressed as the mean ± S.E.M..* p < 0.05.

We then tested for effects of cKO of GR on hippocampal density of OPCs and OLs, stained with NG2 and Aspartoacylase (ASPA, marker for mature myelinating OLs), respectively. While we found no change in the density of OPCs (Fig. 1 *E* and *F*), there was a decrease in the density of mature OLs (Fig. 1 *G* and *H*) in the hippocampus of cKO mice. These data were confirmed by FACS analysis, which showed no difference in the number of PDGFRA+ OPCs and a decrease in CNPase+ OLs in the CD45-CD11b- global populations (Fig. 1 *K* and *L*).

Given an imbalance in OPC and OL ratio may be a consequence of an alteration in OPC proliferation (33), we performed IF staining for Ki-67, a marker for cell proliferation, and PDGFRA, a marker for OPCs, and quantified the overall density of OPCs (PDGFRA+ cells), proliferative cells (Ki-67+ cells) and proliferative OPCs (PDGFRA+ Ki-67+ cells). The density of proliferative OPCs was comparable between cKO and littermate controls (Fig. 1 *I* and *J*), in contrast, a slightly different pattern was observed in the cortex. Indeed, while a reduction of OPC density was observed when employing the NG2 marker (*SI Appendix*, Fig. S2 C), no changes were observed in OL density (*SI Appendix*, Fig. S2 D), OPCs proliferation (*SI Appendix*, Fig. S2 E) or OPC density, when using PDGFRA as a marker (*SI Appendix*, Fig. S2 G). The absence of difference in the density of OPCs and OLs in the cortex was further validated by FACS analysis (*SI Appendix*, Fig. S2 H and I).

A fundamental feature of OPCs is their morphological complexity that allows for continuous and efficient surveillance of the surrounding environment and has been shown to diminish when OPCs undergo apoptosis or differentiation into mature oligodendrocytes (13, 26). Critically, Sholl analysis showed that GR KO in OPCs did not influence filament cell length (*SI Appendix*, Fig. S5 A and B) or complexity of the arborization (*SI Appendix*, Fig. S5 C). Qualitative analysis of single OPC complexity did not suggest heterogeneity within the cell population (Fig. S5 D-E). This indicates that loss of GR is unlikely to severely alter OPCs surveillance capabilities and, in line with the density readouts, does not lead to cell atrophy, such as in response to stressors prior to apoptosis (26).

To assess whether the reduction in mature oligodendrocyte density was reflected in an overall difference in myelin content at ultrastructural level, we performed electron microscopy on hippocampal specimens (Fig. 2 *A*). The relation between g-ratio and axon diameter was estimated in male (Fig. 2 *B*) and female mice (Fig. 2 *C*). While there were small sex-differences in the g-ratio (slightly higher in females) (Fig. 2 *D*), but no genotype or sex differences in axon diameter (Fig. 2 *E*), we found a decrease in the density of myelinated axons in both sexes of cKO mice (Fig. 2 *F*). When analyzing myelin-related proteins, we did not detect any differences in MBP (all isoforms analyzed together), PLP (34), CNP, or ASPA analyzed by western blot (WB), in the hippocampus and cortex of cKO and control mice (*SI Appendix*, Figs. S6 and S7). We also performed 3D reconstruction and morphological analysis of the paranodes, nodes of Ranvier and length of the internodes in control vs. cKO mice and found no differences in sex and genotypes between the groups (*SI Appendix*, Fig. S8).

Overall, these data demonstrate that loss of GR in OPCs in early life reduces the density of oligodendrocytes and of myelinated axons in the hippocampus of adult mice, without affecting myelin ultrastructure. The absence of changes in myelin protein content, axon-myelin units, and overall internode architecture further indicates that GR loss selectively alters specific features of the OL lineage and myelination within the hippocampus.

### Early conditional deletion of GR in OPCs alters the excitability of the neuronal network in response to acute challenge later in life

We tested whether the deletion of GR in OPCs impacts neuronal network functions, using electrophysiological multi-electrode array (MEA) recordings in acute hippocampal slices under control conditions and in the presence of the potent GR agonist dexamethasone (35). This experimental condition mimics *in vitro* the activation of the hypothalamic-pituitary-adrenal (HPA) axis and, thus, the GR-related pathway. Horizontally-sectioned brain slices were placed on the MEA chip to allow measurement of extracellular field excitatory postsynaptic potentials (fEPSPs) along the Schaffer Collateral CA3 to CA1 pathway (Fig. 3 inset; for details see Materials and Methods). Electrically evoked fEPSPs were recorded in the CA1 region, with gradually increasing signal amplitudes plotted for male and female adult mice (Fig. 3 *A* and *B*). Control experiments with 6,7-dinitroquinoxaline-2,3-dione (DNQX) application verified that the evoked signals in CA1 were AMPAR-dominated (*SI Appendix*, Fig. S9). While there were no effects of dexamethasone treatment on the evoked mean fEPSP amplitudes in control and cKO male mice (Fig. 3 *C*), increases were observed in control, but not cKO females for stimulation inputs ranging from 3.5 – 5V (Fig. 3 *D*). The missing increase of the fEPSP amplitude in female NG2-specific GR cKO mice implies that this receptor in OPCs mediates acute neuronal stress responses in females. In addition, the dexamethasone-induced increase of fEPSP amplitudes in female control mice indicated, at least in part, a sex-specific change of the postsynaptic AMPAR-function during acute stress.

**Figure 3.**
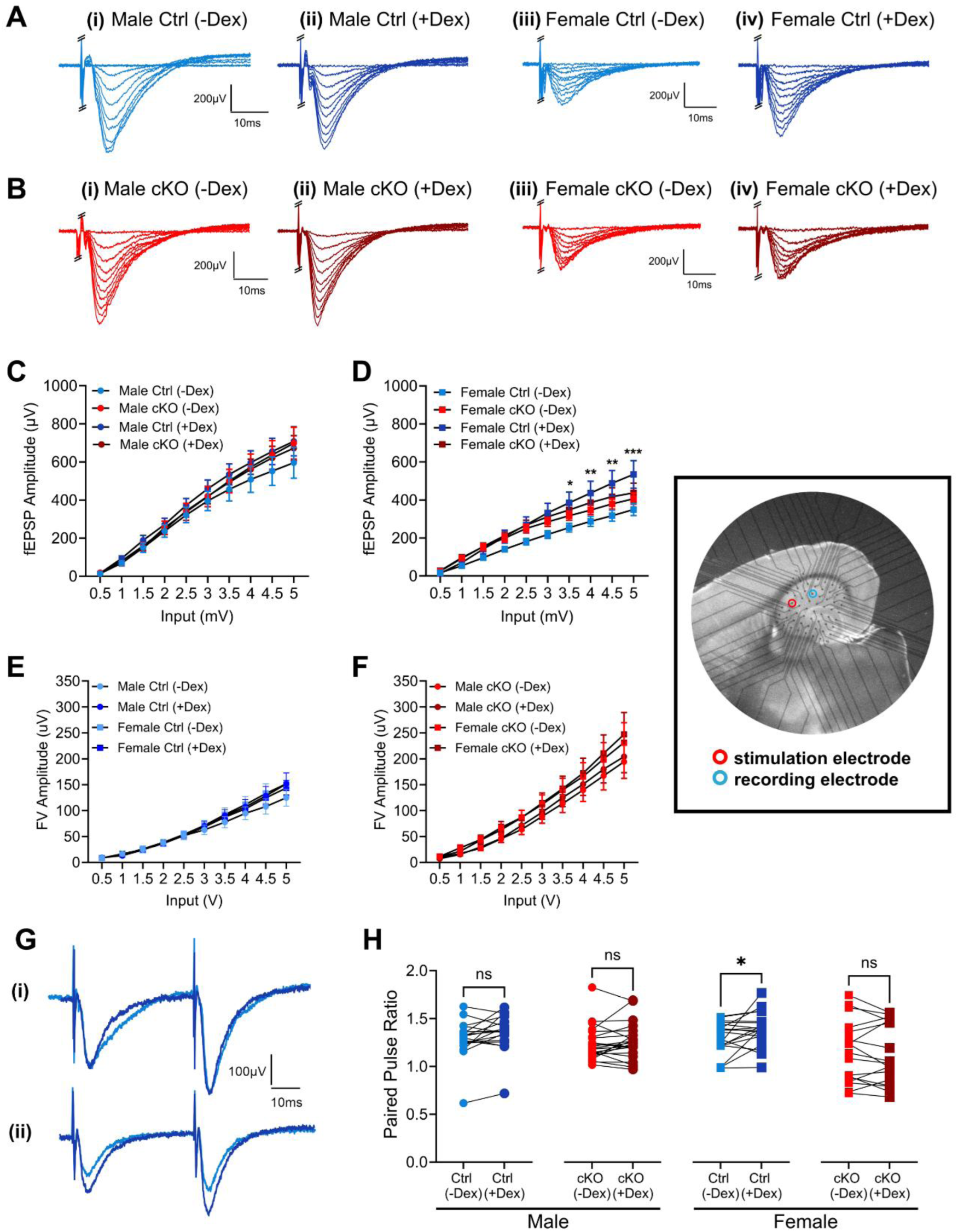
OPCs-specific GR affects hippocampal network excitability under acute stress conditions selectively in adult female mice. (**A**) Representative overlaid voltage traces of the evoked I/O responses from male wildtype mice before (**i**) and after (**ii**) dexamethasone application, and from female wildtype mice before (**iii**) and after (**iv**) dexamethasone application. (**B**) Representative overlaid voltage traces of the evoked I/O responses from male cKO mice before (**i**) and after (**ii**) dexamethasone application, and from female cKO mice before (**iii**) and after (**iv**) dexamethasone application. (**C**) Mean evoked fEPSP amplitudes at different stimulus intensities in the CA1 region of adult male wildtype and cKO mice before (light blue/red, respectively) and 30 minutes after (dark blue/red, respectively) bath application of dexamethasone (**Three-factor statistical model, male**. Three-Way RM ANOVA: input, F(9.000, 342.0) = 166.0, p < 0.0001; n = 18 WT(-Dex), n = 22 cKO (-Dex), n = 18 WT (+Dex), n = 22 KO (+Dex)). (**D**) Same recordings as in (A), but here performed in adult female wildtype and cKO mice (Three-factor statistical model*, female*. Mixed-effects model: Input x Genotype x Dex, F(1.162, 35.88) = 3.622, p = 0.0539; *Statistical analysis per genotype – Female, Ctrl.* Mixed-effects model: Input x Dex, F(1.128, 16.80) = 5.702. Paired t test as post-hoc comparison: input 3.5: (-Dex) < (+Dex), p = 0.0406; input 4: (-Dex) < (+Dex), p = 0.0329; input 4.5: (-Dex) < (+Dex), p = 0.0249; input 5: (-Dex) < (+Dex), p = 0.0238; n = 17 (-Dex), n = 16 (+Dex). *Statistical analysis per genotype – Female, cKO.* Two-way RM ANOVA, Input, F(1.402, 22.43) = 90.15, p < 0.0001; Dex and interaction, p = 0.2581; n = 17 (-Dex), n = 17 (+Dex)). (**E**) Mean fiber volley (FV) amplitudes recorded at different stimulus intensities in the CA1 region of adult male and female wildtype mice before (light blue circles/squares, respectively) and 30 minutes after (dark blue circles/squares, respectively) bath application of dexamethasone (Three-way RM ANOVA, input F(9, 279) = 103.3, p <0.0001; drug, sex p>0.05) (**F**) Same recordings as in (E), but here performed in cKO animals (Three-way RM ANOVA, input F(9, 288) = 76.00, p <0.0001; drug, sex p>0.05). (**G**) **(i)** Representative voltage traces of male Ctrl paired-pulse stimulation responses before (light blue) and after (dark blue) dexamethasone application. **(ii)** Representative voltage traces of female Ctrl paired-pulse stimulation responses before (light blue) and after (dark blue) dexamethasone application. **(H)** Summary plots of the paired-pulse ratio (PPR) of fEPSP-amplitudes recorded in male and female Ctrl- and cKO-mice, before and after 30-minute dexamethasone application (**Female, Ctrl**. Paired t test: t(18) = 2.247, p = 0.0374; n = 19 (-Dex), n = 19 (+Dex). **Female, cKO**. Wilcoxon matched-pairs signed rank test: W = −31.00, p = 0.4874; n = 17 (-Dex), n = 17 (+Dex)). The inset shows a representative acute hippocampal brain slice section on the MEA chip positioned with the stimulating electrode below the Schaffer Collaterals in area CA3 and the recording electrode below area CA1.Data are expressed as the mean ± S.E.M. * p < 0.05

Importantly, we cannot exclude that the sex and/or GR cKO affected the axonal excitability of the presynaptic Schaffer collaterals, so we further analysed the fiber volley (FV) component in our recorded fEPSP signals (Fig. 3 *E*, *F*). Here we failed to detect any significant sex-specific changes in the mean amplitudes of the FV-component before and in presence of dexamethasone treatment (Fig. 3 *E*, *F*). This indicates that axonal excitability does not mediate the observed increase of fEPSP-amplitudes in female mice. Still, slices from GR cKO mice generally reached larger mean FV-amplitudes, linking to an additional contribution of the genotype. This was also visible when plotting the mean fEPSP amplitudes against the FV signal (see *SI Appendix*, Fig. S10). When we further inspected the voltage traces of evoked fEPSPs before and after 30-minute application of dexamethasone, we observed a sex-specific dichotomy that was evident prior to application of dexamethasone, given the higher evoked hippocampal excitability at baseline in both male wildtype and cKO animals as compared to female mice (*SI Appendix*, Fig. S11 A and B). We also performed a paired-pulse stimulation protocol to investigate whether dexamethasone application, mimicking acute stress exposure, induces changes in short-term synaptic plasticity at the Schaffer collateral pathway (Fig. 3 *G*). Dexamethasone application caused a larger increase in the paired-pulse ratio (PPR) only in the female control mice, whereas no effects were recorded in male control mice, cKO mice and in female cKO mice (Fig. 3 *H*). These results reflect those from the evoked fEPSP experiments above (Fig. 3 *C, D*). Representative traces of the paired-pulse stimulation responses, with a 50ms interstimulus interval, show a lack of change in amplitude of either the first or second paired-pulse stimulation responses following application of dexamethasone for male controls (Fig. 3 *G* (i)). In contrast, an increase in the amplitude of both the first and second paired-pulse stimulation responses following dexamethasone application was observed in female controls, with the second stimulation undergoing a relatively larger increase (Fig. 3 *G* (ii)). The changes in paired-pulse stimulation responses indicated an altered synaptic facilitation, in which the probability of neurotransmitter release by the second stimulation remains high (Fig. 3 *H*). We also found an overall slight genotype difference in PPR prior to dexamethasone application (*SI Appendix,* Fig. S11 C, see *SI Appendix,* Table S1 for the detailed Statistics). We found no changes in the rate of spontaneous network activity in the hippocampal slices of either sex or genotype after dexamethasone application (*SI Appendix*, Fig. S11 D). The neuronal spontaneous activity in these MEA recordings was verified by application of Tetrodotoxin (TTX), which blocked all signs of spontaneous activity (*SI Appendix*, Fig. S9). A sex-specific dichotomy was again observed in the mean spontaneous spike frequency before dexamethasone was applied, with female mice tending to have a higher mean spike frequency than male mice (*SI Appendix*, Fig. S10 E).

In summary, the electrophysiological experiments suggest that acute stress primarily affects the hippocampal network in female control mice, divulging a novel and fundamental sex-specific function of the NG2-cell-specific GR in the hippocampal Schaffer collateral pathway.

### Altered long-term potentiation in acutely stressed female cKO mice

As the electrophysiological experiments indicated a relevance of the NG2-GR for acute stress responses in female mice, female wildtype and cKO mice were then further tested for alterations in long-term potentiation (LTP) (Fig. 4 *A*). We found that both wildtype and cKO mouse lines were capable of LTP induction and maintenance in the presence of bath-applied dexamethasone that mimics an acute *in vivo* stressor in our *in vitro* slice preparation (Fig. 4 *A*). However, LTP in cKO slices exposed to dexamethasone was lower than in wildtype slices (Fig. 4 *B* and *C*). Thus, the cKO of GR in NG2 cells impairs hippocampal LTP along the Schaffer Collateral pathway, from the CA3 to the CA1 region, where reduced LTP may indicate impairments in behavioral tasks assessing learning and memory in female cKO mice under acute stress.

**Figure 4.**
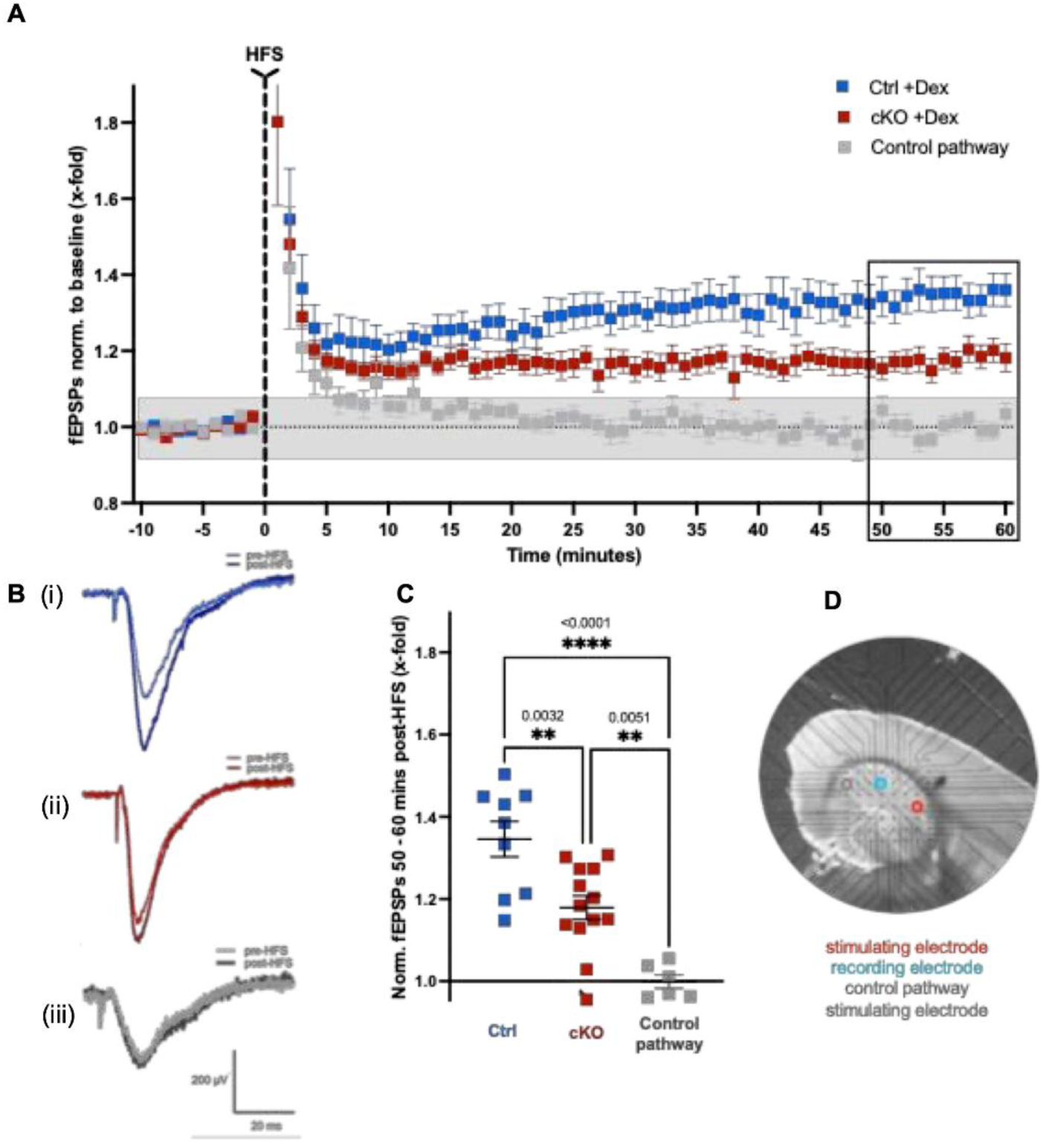
Female NG2-cKO mice show an impaired hippocampal LTP compared to female control mice. **(A)** LTP was induced at the Schaffer collaterals using a 100 Hz high-frequency stimulation (HFS) protocol and recorded in area CA1 using a microelectrode array (MEA) in female control and female NG2 cKO mice. Blue points represent the control female mice, red points represent the NG2 cKO female mice, and the grey dots represent the recordings taken from the independent control pathway. Graph shows mean +/- S.E.M. of the relative strength of potentiation of fEPSPs. Female control n = 9 slices from 3 mice, female cKO n = 13 slices from 5 mice, independent control pathway recordings n = 6 slices from 3 mice. **(B)** Representative fEPSP traces from both female control (blue) and female cKO (red) genotypes along with traces from the independent control pathway (grey). The darker of each colour shows fEPSPs before the 100Hz stimulation, the lighter of each colour shows fEPSPs 60 minutes after the stimulation. **(C)** The relative strength of potentiation of the fEPSPs recorded at 50 to 60 minutes after the 100Hz stimulation shows that both female genotypes had an LTP that was significantly greater than the control pathway (One-way ANOVA, F(2,25) = 19, p < 0.0001. Tukey’s multiple comparison test: female control > control pathway, p < 0.0001, female cKO > control pathway, p = 0.0051). However, an impairment of LTP in the female cKO genotype compared to the female control genotype can be seen from their significantly lower fEPSP potentiation post-HFS (One-way ANOVA, F(2,25) = 19, p < 0.0001. Tukey’s multiple comparison test: female Ctrl vs female cKO, p = 0.0032). **(D)** Representative hippocampal section with an example of the location of stimulating electrode (red), recording electrode (blue) and stimulating electrode of the independent control pathway (grey) along the Schaffer Collaterals. Statistical analysis using one-way ANOVA with post-hoc Tukey’s multiple comparisons test. Graph shows mean +/- S.E.M.

### Postnatal deletion of GR in OPCs induces impairment in memory and aversive learning tasks but does not alter anxiety-relevant behavior or sociability in adulthood

Given changes in neuronal network activity and LTP may have consequences for behavioral outcomes and are linked with the processes of learning and memory formation (36), we performed a battery of tests encompassing a variety of behavioral domains in control littermate and cKO mice. We tested mice from the less to the more aversive tests and performed an open field (OF) test to assess locomotor activity, novel object recognition test (NORT) for cognitive performance, three-chamber sociability test for sociability, dark-light box (LB) for anxiety-relevant behavior, ending with a two-way active avoidance test (TWA) for aversive learning. When mice were tested in the OF arena, there were no genotype differences in locomotor activity (Fig. 5 *A*), neither in the time spent in the more anxiogenic region of the arena at the center (Fig. 5 *B*) nor at the lower anxiogenic region of the arena at the periphery (rim) (Fig. 5 *C*); however, females of both genotypes spent more time in the lower anxiogenic part of the OF arena (see *SI Appendix,* Table S1 for the Statistics). For assessment of cognitive performance, we first checked in the training phase of the NORT that the animals, which were randomly assigned to an identical object pair, did not show a preference for one of the two identical object positions (Fig. 5 *D*). After an inter-trial time of 24h, one of the two objects was replaced and performance in novel object recognition was calculated and found to be lower in both male and female cKO mice than the control littermate (Fig. 5 *E*). Additional automatically annotated motion tracking analyses via DeepLabCut (37) revealed a general sex difference in mouse positioning relative to the objects over time (*SI Appendix,* Fig. S12). Then we found that neither genotype nor sex showed differences in preference towards a social target in the three-chamber sociability test (Fig. 5 *F*) and in the time spent in the lit compartment of the LB (Fig. 5 *G*), where there was equal first latency to the dark compartment (Fig. 5 *H*) and a similar number of lit-dark compartment transitions in the LB test (Fig. 5 *I*). Finally, both control and cKO mice in the TWA test learned the task over time and increased progressively the number of conditioned responses (i.e. to avoid foot-shock upon a tone presentation) (Fig. 5 *J*), with an interaction between day of the test, sex and genotype. To explain the roles of sex and genotype in this aversive learning test, we separated the learning curves of males (Fig. 5 *K*) from females (Fig. 5 *l*) and found that learning was poorer on day 2 in female cKO mice, which needed more trials to learn the task compared to the control littermates (Fig. 5 *l*). Furthermore, when looking at the daily learning curves we found additional differences. The performance of both male and female cKO mice was worse on the second day of the test (*SI Appendix,* Fig. S13) and they were thus slower in learning the task. Importantly, these cognitive alterations were specific to GR deletion during postnatal development and were not observed following GR deletion in adulthood (*SI Appendix,* Fig. S14). The efficiency of the cKO in adulthood was confirmed by IF showing a reduction of GR in OPC (*SI Appendix,* Fig. S15). To control for the effects of tamoxifen injection, a batch of mice without tamoxifen-induced GR deletion was subjected to the same behavioral tests and showed no behavioral alterations (*SI Appendix,* Fig. S16).

**Figure 5.**
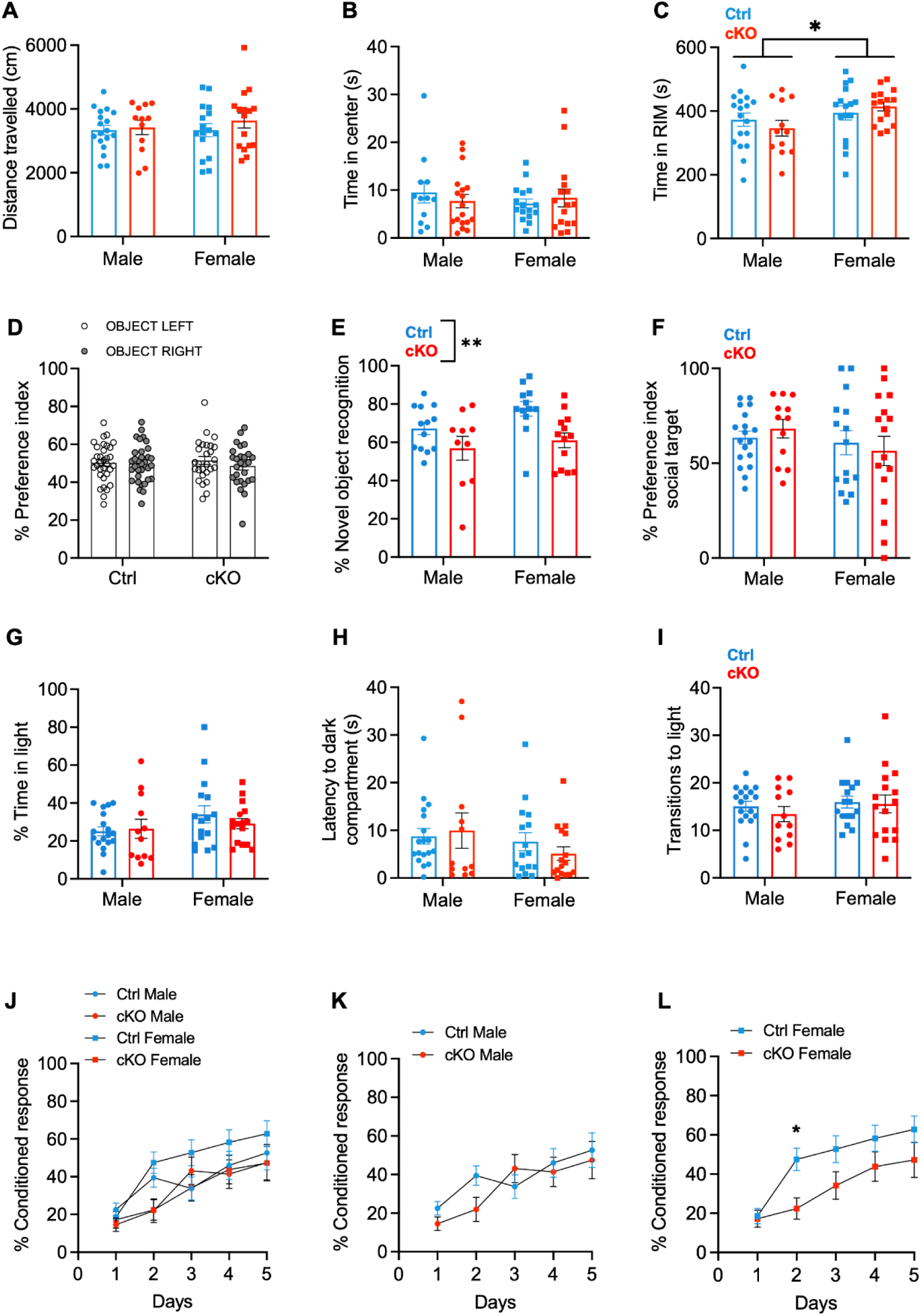
GR deletion in OPCs induces impairment in memory and aversive learning tasks. (**A-C**) **Open Field Test.** (**A**) Distance travelled (Two-way ANOVA: p = 0.2718). (**B**) Time spent in the center (Two-way ANOVA: p = 0.605). (**C**) Time spent in the RIM (Two-way ANOVA: sex, F(1, 58) = 4.707, p = 0.0342; n = 18F/16F Ctrl and n = 12M/16F cKO). (**D-E**) **Novel object recognition test.** (**D**) Preference index (%) for two identical objects (Two-way ANOVA: p = 0.3910), and (**E**) novel object recognition (%) (Two-way ANOVA: genotype, F(1, 44) = 10.02, p = 0.0028; n = 13M/12F Ctrl and n = 10M/13F cKO). (**F**) Sociability index (Two-way ANOVA: p = 0.229; n = 18M/16F Ctrl and n = 12M/16F cKO). (**G-I**) **Light-Dark Box Test.** (**G**) Time spent in the lit compartment (Two-way ANOVA: p = 0.1177). (**H**) Latency to enter the dark compartment (Two-way ANOVA: p = 0.1677). (**I**) Frequency of light-dark transitions (Two-way ANOVA: p = 0.3087; n = 18M/16F Ctrl and n = 12M/16F cKO). (**J-L**) **Two-way active avoidance test.** (**J**) Percentage of conditioned response (CR) over days in both sexes (Three-way ANOVA: genotype x day x sex, F (4, 224) = 2.555, p = 0.0398; n = 17M/16F Ctrl and n = 12M/15F cKO). (**K**) Percentage of conditioned response (CR) over days in males (Two-way RM ANOVA: days x genotype, F(4, 108) = 2.664, p = 0.0363; Šídák’s multiple comparisons test: day 2: cKO = Ctrl). (**L**) Percentage of conditioned response (CR) over days in females (Two-way RM ANOVA: days x genotype, F(4, 116) = 3.010, p = 0.021; Šídák’s multiple comparisons test: day 2: cKO < Ctrl, p = 0.0339; n = 17M/16F Ctrl and n = 12M/15F cKO. Data are expressed as the mean ± S.E.M. *P < 0.05, **P < 0.01.

These data demonstrate that loss of GR in OPCs in early life does not impact on anxiety-relevant behaviors, but leads to deficits in hippocampus-dependent non-aversive and aversive tasks for learning and memory in adulthood, under moderate stress activation of the HPA axis (38–41).

## Discussion

We established here a novel physiological role of GR-signaling in OPCs during postnatal brain development. Specifically, our findings demonstrate that early postnatal deletion of the GR in OPCs influences the density of hippocampal OLs and of myelinated axons in adulthood, alters neuronal network excitability and synaptic potentiation in response to dexamethasone challenge and affects aversive learning later in life. Collectively, our data highlight that constitutive expression of GR in OPCs is critical for maintaining a physiological OPCs-OLs ratio and for supporting hippocampal activity and plasticity and ensuring hippocampus-related cognitive function.

First, we demonstrated that the cKO of GR in postnatal OPCs alters the density of OLs in the adult hippocampus, without affecting the density and proliferation of OPCs (Fig. 1). Previous studies have examined the role of GCs on the proliferation-maturation dynamics of the OLs in murine models. For instance, the exposure to ELA, which activates the HPA axis and elicits an increase in GCs (24, 42), leads to precocious OLs differentiation in pups, and is associated with a depletion of the OPCs pool in adult mice (22). These findings have also been corroborated by post-mortem studies from human individuals with a history of childhood trauma (23). In contrast, in our experimental design, mice were not exposed to stress, and we focused instead on the physiological function of GCs by conditionally deleting GRs in OPCs during the early postnatal period. Our data suggest that physiological GC levels are essential for proper maturation of OPCs into OLs. Furthermore, GCs appear to play a key role in coordinating the timing of differentiation, as previously shown *in vitro*, via signaling clocks in precursor cells during development (43). In support of this role of GCs *in vitro* studies have shown differences in OPC proliferation and differentiation in response to corticosterone treatment (44) and reduction in myelination in mixed central nervous system cell culture after GC treatment (45). In our study, we did not detect changes in myelin ultrastructure and thickness or internode length (Fig. 2; *SI Appendix*, Fig. S8), but we observed a decreased density of myelinated axons in both male and female cKO mice (Fig. 2). A reduced density of myelinated axons has previously been reported in different models, including in post-mortem tissue of patients with progressive multiple sclerosis but in normal-appearing white matter (46). It is tempting to speculate that the decrease in oligodendrocyte density is linked to the decrease in myelinated axon density, with consequences on the development of neuronal circuits.

While our approach, to the best of our knowledge, represents the first mouse model in which GRs have been cKO specifically in OPCs (Fig. 1), with an absence of recombination in other GR-expressing cell types in the CNS (*SI Appendix*, Fig. S4), off-target recombination has been reported for this inducible Cre line in other cell types (47). Therefore, our results may depend on limited off-target recombination in other cells that express NG2 and GR.

Postnatal GC concentrations are tightly controlled to shield the developing brain from stress-induced GC surges, a phenomenon known as the SHRP, during which moderate stressful challenges fail to trigger a significant stress hormone response (2). Our data highlight a critical role for GR signaling in OPCs during this postnatal period, when basal levels of GCs are extremely low. We conclude that postnatal OPCs depend on a delicate regulation of circulating GCs. Elevated levels of GCs, as observed during ELA, may disrupt OL lineage dynamics, resulting in changes associated with negative health outcomes. Conversely, maintaining homeostatic levels of GCs is likely crucial for supporting proper oligodendrocyte lineage maturation.

Beyond the canonical role of OPCs in sustaining oligodendrogenesis and myelination of axons, our data also revealed a sex-specific role of GR in OPCs in modulating neuronal network excitability and potentiation upon challenge. We demonstrated that acute application of the synthetic GC agonist, dexamethasone, increased fEPSPs, PPR in control female mice, but not in cKO mice. We also showed that female cKO has a lower LTP compared to control after incubation with dexamethasone. These data raised two important new points: first, they revealed a clear sex-specific electrophysiological response under acute challenging conditions (i.e. *ex vivo* dexamethasone application), indicating that female mice might be more sensitive to slight increases in circulating stress hormone and potentially more responsive to HPA activity (48–51). Second, GR in OPCs appears to be necessary to induce changes in neuronal network excitability and synaptic potentiation in response to a mild and/or acute stressor. It has been shown previously that GCs modulate learning and memory processes. GCs also modulate synaptic transmission by slowly enhancing the amplitude of miniature excitatory postsynaptic currents (mEPSCs) (52) and rapidly facilitating synaptic potentiation in the mouse hippocampal CA1 area (53); however, these data were always obtained from patch-clamped neurons. Here, we have shown for the first time that a specific non-neuronal cell type is required for such excitability and it is mechanistically involved in synaptic plasticity, which may be relevant for memory and learning. Whether this phenomenon is mostly driven by the dorsal or ventral region of the hippocampus remains an open question that cannot be addressed through our dataset. Although dexamethasone is a potent GR agonist, it can also bind to mineralocorticoid receptors (MR) with low affinity. In our study, we did not perform additional experiments in the presence of a GR antagonist such as RU486, therefore, the possibility that MR receptors in neighboring neurons may also be activated in cKO mice as compensatory effects, due to the lack of GR in OPCs cannot be completely ruled out. However, previous studies have shown that dexamethasone at the dose used in our study induces LTP-related mechanisms that are completely blocked by RU486 (54), supporting a predominant role of GR in mediating these plasticity effects. Based on our data that showed changes in the density of myelinated axons and differences in network excitability and synaptic plasticity in response to dexamethasone, we speculate that postnatal deletion of GR in OPCs may have long-lasting consequences for neuronal network formation. Rechallenging such neuronal networks in the presence of dexamethasone is also impaired, possibly indicating a direct OPC–neuron interaction, conceivably involving GR signaling in OPCs.

The sex- and brain region-specific alterations observed *ex vivo* in the GR cKO mice in network excitability and LTP occurred coincidently with an impaired performance in the non-aversive memory test (NORT) and in an aversive learning paradigm (TWA), especially in female mice. In the NORT, mice are exposed to novelty, which can be arousing, but is unlikely to lead to substantial increase of circulating GCs (55). In the TWA, mice are exposed to a mild stressful situation which triggers the activation of the HPA axis (41). It has previously been shown that fear conditioning induces OPCs to proliferate and differentiate into myelinating oligodendrocytes (56) and that myelin formation is necessary for remote fear memory retrieval. Similar data were obtained for non-aversive learning, including motor learning or hippocampal-related learning and memory in the Morris water maze (57). Clinical data also support the importance of experience-dependent myelin formation. Indeed, structural MRI studies have shown that white matter changes are required for the acquisition of both new motor and cognitive skills (e.g. learning to read, juggling, piano playing) (58–60). Moreover, cognitive deficits have been associated with (dys)function of the oligodendrocyte lineage and alterations in myelin content and ultrastructure. Our cKO mice, which have a reduced density of OLs and of myelinated axons and impairment in LTP, showed deficits in the NORT and were slower than the controls in learning the task in the TWA. In these mice, the reduction in mature OLs was associated with a decrease in the density of myelinated axons, however the myelinated axons showed normal myelin content and g-ratio. This suggests that the extant OLs, despite being reduced in the cKO mice, may not be sufficient to myelinate all hippocampal axons. These changes in neuronal circuitries could impair baseline cognitive performance, such as those observed in the NORT, and may also affect the excitability and plasticity of the circuitry in response to dexamethasone treatment and of the behaviors of the mice under mild stressful situations, such those observed in the TWA.

GCs play a critical role in memory formation (61, 62): both memory-enhancing and -impairing effects have been reported (61, 63). Extending the classical view that GCs exert their cognition-modulating effects exclusively by modulating neuronal function, a recent study showed that the astrocyte-specific ablation of GR led to impaired aversive memory expression in two different paradigms of Pavlovian learning (64). Thus, it is tempting to speculate that proper memory formation, initiated by activation of the HPA axis, requires GR expression in OPCs during development. Specifically, GR signaling in OPCs may regulate the proper formation of neuronal circuitry and its capacity to respond to acute challenges later in life; disruption of this process could impair OPC-neuron communication with consequences on memory formation and learning in adulthood.

In conclusion, our data demonstrate that GRs are essential from the very early stages of postnatal development to regulate OL differentiation and axon myelination. While the specific developmental check-points of the GC pathway in OPC remain unexplored, we demonstrated that the loss of GRs in early life OPCs appears to prime hippocampal network excitability and plasticity and impair learning abilities, particularly under mild stress in adulthood. Based on our histological findings, it is apparent that the lack of GR in OPCs impacts on their canonical, myelin-related functions. However, the reduced responsiveness of the hippocampal network to dexamethasone application in cKO suggests that the GC pathway in OPCs might also be essential for non-canonical, myelin-independent regulation of network excitability. Future studies are needed to further clarify the conditions regulating each of these functions and how they orchestrate cognitive processes in response to environmental challenges.

## Materials and Methods

A succinct description of the experimental procedures related to the main results is provided below. Comprehensive materials and methods descriptions, including methods related to supplementary results, are reported in the *SI Appendix,* Supplementary Material and Methods.

### Mice

NG2-CreER^T2^ male mice (B6.Cspg4^tm1.1(cre/ERT2)Fki^) (47) were crossed with female Nr3c1^fl/fl^ (B6.Cg-Nr3c1^tm1.1Jda/J^, Jacksons Lab, #021021) to obtain NG2-CreER^T2^ x Nr3c1^fl/fl^ mice (cKO mice), which expressed tamoxifen-dependent Cre recombinase under the control of the NG2 transcriptional regulatory elements and Nr3c1 gene flanked by two loxP sites. NG2-CreER^T2^ x Nr3c1^fl/fl^ male mice were then bred with Nr3c1^fl/fl^ female mice to obtain the experimental mice. Littermate Nr3c1^fl/fl^ were employed as control (Ctrl).

#### Early postnatal deletion cohort

Tamoxifen (TAM, Sigma-Aldrich, T5648) was dissolved in pure ethanol (Laborchem® international, LC-8657.1) and diluted in corn oil (Sigma-Aldrich, C8267) at a working concentration of 20 mg ml^-1^. All the pups included in the experiment received i.p. injection of 0.1 mg tamoxifen in 5 μl vehicle at PDs 2 and PD 4 (65). The mice were allowed to reach adulthood undisturbed, except for the conventional caretaking routine procedure in the mouse unit.

#### Adult deletion cohort

Tamoxifen (TAM, Sigma-Aldrich, T5648) was dissolved in pure ethanol (Laborchem® international, LC-8657.1) and diluted in corn oil (Sigma-Aldrich, C8267) at a working concentration of 20 mg ml^-1^. All the young adult (> P60) mice included in the experiment received 75 mg/kg TAM daily for 5 consecutive days via i.p. injection. Body weight and physical status were monitored throughout. The mice were left undisturbed for additional 3 weeks, to allow for recombination and complete depletion of the endogenous GR.

#### Mouse line control cohort

To control for any confounding effects from the lack of one NG2 allele, independently of the deletion of GR, and to validate the selected control group, male and female adult (>10-week-old) NG2-CreER^T2^ x Nr3c1^fl/fl^ and Nr3c1^fl/fl^ mice underwent physical and behavioral assessment, without TAM treatment.

Genotyping was performed as described before (47) or adapted from the recommended protocol on the JAX datasheet via standard PCR of genomic DNA. Primer sequences are reported in the supplementary materials and representative genotyping output in *SI Appendix,* Fig. S2 A. All experiments were then conducted on adult mice of both sexes (8-12 weeks old). Mice of the same sex were grouped housed (3-5 per cage) and maintained at a 12/12h light-dark cycle (lights on at 7:00 A.M.), at controlled temperature and humidity (temperature = 22 ± 2 ^◦^C, relative humidity = 50 ± 5%). Food and water were available *ad libitum*. All experiments were performed in accordance with the European directive 2010/63/EU for animal experiments and were approved by the local authorities (Animal Protection Committee of the State Government, Landesuntersuchungsamt Rheinland-Pfalz, Koblenz, Germany).

### Flow cytometry

Naïve mice were sacrificed with isoflurane (Piramal Critical Care Deutschland; Cat# 4150097146757) and perfused with ice-cold 0.1 M pH 7.4 phosphate buffer saline (PBS) (ThermoFisher Scientific, 10010023). The hippocampi and cortices were rapidly dissected and coarsely chopped in Hank’s Balanced salt solution (HBSS + CaCl_2_, MgCl_2_) containing 1 mg/ml of papain (from papaya latex, dnase I Sigma-Aldrich, 51807363), including 40 μg/ml of DNase I (grade II from bovine pancreas, Sigma-Aldrich, 10104159001 (51743558)). Tissue was dissociated using gentleMACS™ Octo Dissociator (Miltenyi) using a built-in ready-to-use program optimized for small quantities (<100 mg) of the CNS tissue. The cells were then resuspended in 8 ml DMEM/ 1% horse serum and filtered through 70-μm cell strainers. The resulting cell pellet was then re-suspended in 25% Percoll solution (Cytiva) and centrifuged (25 min, room temperature (RT), 300 *g*). Cells were resuspended in MACS Neuro Medium (Miltenyi Biotec, 130-093-570) containing 1x MACS NeuroBrew-21 (Miltenyi, 130-093-566), 100 U/ml Penicillin Streptomycin (Thermo Fisher Scientific, 15140122) and 2 mM L-Glutamine (Thermo Fisher Scientific, 25030-024 (51709381), and incubated with slow rotation at 37 °C for two hours. After washing the cells in FACS buffer (0,5% bovine serum albumin (PAN-Biotech P06-1391500 (51809570)), 2 mM Ethylenediaminetetraacetic Acid (EDTA, Sigma-Aldrich) in PBS), Fc receptors were blocked for 10 min with Fc-block (1:100 in FACS buffer, BioXCell, BE0307). Surface antigens were then labelled for 20 min on ice with FACS buffer containing following reagents: 1:1000 Fixable Viability Dye eFluor™ 780 (eBioscience, 65-0865), PE/Cyanine7 anti-mouse/human CD11b 1:1000 (BioLegend 101216), Brilliant Violet 510™ anti-mouse CD45 (BioLegend, 103138) and CD140a (PDGFRA) APC 1:10 (Miltenyi Biotec, 130-102-473). For intracellular staining cells were fixed and permeabilized with eBioscience™ Foxp3/transcription factor staining buffer set (ThermoFisher Scientific, 00552300) according to the manufacturer’s instructions. Cells were analyzed by using a FACSCanto II device (BD Biosciences). Post-acquisition analysis was performed by using FlowJo software. A list of the employed antibodies is provided in the supplementary material.

### Immunohistochemistry

Mice were anesthetized via i.p. of Ketamin/Xylaxin (200 mg/kg, 20 mg/kg) and intracardially perfused with ice-cold PBS followed by 4% paraformaldehyde (ChemCruz). Brains were post-fixed in 4% PFA at 4°C for 2 hours, with the exception of Pdgfra and Ki-67 staining and for Caspr, Mbp and Nav1.6 (post-fixed 24 hours), then transferred into 30% sucrose in 0.1 M PBS, pH 7.4 for 48 h and afterwards embedded in a cryoprotective medium (Tissue-Tek, O.C.T., 4583, Sakura Finetek Europe). Free-floating coronal sections (30-35 μm) including the somatosensory cortex and dorsal hippocampus (bregma from −1.79mm to −2.53mm) were washed in PBS (3x20min), then incubated for 90 minutes in 0.5% (v/v) Triton X-100 in, followed by 1 hour of blocking and permeability with 10% goat or donkey serum in 0.5% Triton X-100 in PBS at RT. Pdgfra and Ki-67 and Mbp/Caspr1 immunostaining slices were permeabilized and blocked for 60 min and 150 minutes, respectively. The sections were then incubated overnight (ON) at 4°C in primary antibody, diluted in 3% serum in 0.5% Triton X-100 in PBS. On the following day, sections were washed in PBS (3x20 min) and incubated for 2 hours at RT with secondary antibody diluted in 3% serum in 0.5% Triton X-100 in PBS. The sections were then washed in PBS (3 x 20 min), counterstained with DAPI (1:10000 in ddH20, SIGMA, D9542) and mounted on SuperFrost slides (Epredia, K5800AMNZ72) with fluorescence mounting medium (Dako, S3023). For the paranodes and nodes of Ranvier immunohistochemistry, free-floating coronal sections (30-35 μm) including the somatosensory cortex and dorsal hippocampus (bregma from −1.79 mm to −2.53 mm) were washed in PBS (3x10 min), blocked in 5% NGS, 2,5% BSA, 0,25% TritonX-100 in PBS for 1 hour at RT and incubated ON at 4°C in primary antibody diluted in 5%NGS, 2,5% BSA, 0,25% TritonX-100 in PBS. On the following day, sections were washed in PBS (4x10 min) and incubated for 2 hours at RT with secondary antibodies diluted in 5%NGS, 2,5% BSA, 0,25% TritonX-100 in PBS. The sections were then washed in PBS (4 x 10 min) and mounted on slides with Vectashield plus DAPI (4’,6-diamidino-2-phenylindole) mounting medium (#H-1200, Vector Laboratories). The used antibodies were: rabbit anti-Nav1.6 (1:250, #ASC-009, Alomone labs), mouse anti-CASPR (1:1000, #75-001, Neuromab), goat anti-rabbit Alexa Fluor 594 (1:500, 111-585-003, Jackson Immuno Research), donkey anti-mouse Alexa

Fluor 488 (1: 500, 715-545-150, Jackson Immuno Research). A list of the employed antibodies is provided in the supplementary material. Naïve mice were employed for proliferation and PDGFRA+ cells density assessment and for the analysis of paranodes and nodes of Ranvier length; the remaining samples were collected from mice that underwent behavioral testing (see Fig. 1 *A*).

### Microscopy and image analysis

Images were acquired at a Leica TCS SP8 confocal laser scanning microscope. For cell quantification (density or percentage), confocal micrographs were acquired with a 20x/0.75 (HC PL APO IMM CORR CS2) or 40x/1.30 (HC PL APO OIL CS2) immersion objectives, at a format of 1024 × 1024 pixels. The pinhole was set to 1 AU, the field of view ranged from 276.09 μm × 276.09 μm to 87.87 μm × 87.87 μm, and the voxel size from 270x 270 x 1000 nm^3^ (xyz) to 637nm x 637nm x 1000 nm^3^ (xyz). For each mouse, 4-6 micrographs (2-3 brain sections) from the dorsal CA1 and L2/3 of the somatosensory cortex, were acquired and Z-stack images were processed and analyzed with ImageJ (NIH) software by a blinded experimenter (66). Recombination efficiency was calculated as the proportion of AN2^+^Olig2^+^ cells co-expressing GR. Recombination specificity was estimated based on the proportion of neurons (NeuN^+^), microglia (Iba1^+^), astrocytes (GFAP^+^), and the area covered by pericytes (PDGFRB^+^) co-expressing GR. Cell density was calculated as the number of cells per unit area (cells/mm²) by dividing the total cell count by the area of the region of interest (ROI). OPCs proliferation was calculated as the percentage of PDGFRA^+^ cells co-expressing Ki-67^+^. Only cells with a detectable DAPI signal were included in the analysis. For OPCs morphology analysis, confocal micrographs were acquired with a HC PL APO CS2 63x/1.40 oil objective; pinhole was set to 1AU, the image format to 1024 x 1024 pixel, the pixel size to 86nm x 86nm and the z-stack interval to 150nm. A detailed description of the workflow and analysis parameters are provided in the supplementary material.

### Electron microscopy

Freshly harvested brains were cut into ∼1.5 mm thick slices and fixed in a solution containing 2.5% glutaraldehyde, 1.25% paraformaldehyde, 0.05% picric acid in 0.1 M cacodylate buffer pH 7.2, for a minimum of 24h and then trimmed into smaller areas containing the region of interest (hippocampus with cornu ammonis), washed in a 0.1 M cacodylate buffer and then afterwards post-fixed in 2% Osmium tetroxide, dehydrated with ethanol/propylene oxide, and embedded in glycidyl ether 100 (EPON 812, SERVA Electrophoresis GmbH, Heidelberg, Germany). 0.5 μm semi-thin sections were collected, stained with toluidine blue pyronin and trimmed again in smaller regions. The EPON blocks were then cut using a Reichert Ultracut S ultramicrotome (Leica Microsystems, Wetzlar, Germany) and ultrathin sections (60 nm) were mounted on copper grids, contrasted with 4% uranyl acetate and lead citrate to confirm the brain region and observed with a Zeiss EM 10 C TEM (Carl Zeiss GmbH, Jena, Germany). Images were taken with ImageSP System (SYSPROG, Minsk, Belarus) from each section with a scale of 4.92 nm/ pixel. Myelin thickness (g-ratio), axon diameter and myelinated axon density were quantified manually using FIJI software. The myelin g-ratio was calculated as the ratio of the inner axon diameter (including non-compacting myelin)/ outer axon diameter (including the myelin sheath). For each image, all g-ratios were averaged, and these image means were subsequently averaged across all images collected from a given animal. All the axons contained in the acquired micrographs – total ≥100 axons/region/mouse - were quantified, except for any myelinated axons that presented obvious fixation artefacts or did not have a coronal cut, which were excluded from the analysis.

### Electrophysiological Recordings – Multielectrode Array

Naïve mice of 9-12 weeks, which received TAM at PDs 2 and 4, were used for electrophysiological experiments. After deep anaesthesia using i.p. injection of 200 mg/kg ketamine ketamine, choline-based artificial cerebrospinal fluid (c-ACSF) solution was perfused by injection into one heart ventricle, containing in mM: NaCl (87), KCl (2.5), NaH_2_PO4+H2O (1.25), choline chloride (37.5), NaHCO3 (25) and d-glucose (25), CaCl *2H_2_O (0.5), MgCl_2_ (7) (pH of 7.4). The brain was removed and placed into ice-cold c-ACSF perfused with carbogen (95% oxygen and 5% carbon dioxide), then placed on a vibratome stage to cut horizontal slices with thickness of 350μm. Slices were collected in a storage chamber of normal ACSF, which contains (in mM) NaCl (125), KCl (3), MgSO4 x 7H2O (1.3), CaCl2 x H2O (2.5), NaH2PO4 x H2O (1.25), NaHCO3 (26) and d-glucose (13), to rest at room temperature for at least 40 minutes before use. On the multi-electrode array (MEA) chip (60MEA200/30iR; MEA2100 System, Multi Channel Systems MCS GmbH), the slice was constantly perfused with carbogenated normal ACSF, which also contained 0.01% EtOH in order to control for any ethanol-induced electrophysiological changes (see below). Each slice was precisely placed to position a stimulating electrode in the CA3 region and recording electrode in the CA1 region of the hippocampus, along the Schaffer Collateral pathway. A platinum grid was placed on top of the slice to ensure electrode-tissue contact before starting the recordings. All recording protocols were generated and applied by Multi Channel Experimenter 2.2 software. Spontaneous neuronal activity was measured by recording extracellular spike frequency whereby an event counted as a spike exceeding the downward threshold of five standard deviations from baseline activity. Events were recorded for 5 minutes and quantified by the Multi Channel Analyzer 2.2. For evoked activity and FV analysis, an input-output curve was generated by applying step-wise voltage stimulations that ranged from 0.5V to 5V in 500mV steps. The maximum amplitude of both the FV and each evoked fEPSP was recorded by the electrode in the CA1 region for each slice. The paired-pulse stimulation protocol, which had a 50ms inter-stimulus interval, used the voltage stimulation intensity which produced an fEPSP equal to ∼30% of the maximum evoked fEPSP amplitude from the input/output curve protocol, for each slice. After, the perfusion input was switched to carbogenated ACSF containing 200nM of Dexamethasone diluted in ethanol (0.01%). After wash-in of 30 minutes all protocols were repeated. On finishing, the slice was removed and the drug-infused ACSF completely washed out, before placing the next slice. To verify that fEPSPs were AMPAR-dependant, 6,7-dinitroquinoxaline-2,3-dione (DNQX) at a concentration of 10µM was washed onto the slice for 10 minutes, resulting in the elimination of the fEPSP. To then verify that detected spontaneous spiking activity relied on sodium channel function and not electrical noise or artefacts, tetrodotoxin (TTX) at a concentration of 1µM was washed on for a further 10 minutes, resulting in a silencing of spiking activity.

For LTP experiments, female control (N = 3) and female cKO (N = 5) mice of 9-12 weeks were anaesthetised and the brain removed, sliced and incubated in the same way as above stated. 200nM of Dexamethasone diluted in ethanol (0.01%) was consistently present in the ACSF perfusion of the slices. Again, a voltage stimulation intensity which produced an fEPSP equal to ∼30% of the maximum evoked fEPSP amplitude from the input/output curve protocol was used in the CA3 region, for each slice. After 10 minutes of baseline recordings in the CA1 region, with an interstimulus interval of 60 seconds in the CA3 region, a 100 Hz high frequency stimulation (HFS) was given to induce LTP. This was followed by 60 minutes of baseline intensity stimulations, every 60 seconds. The level of HFS- induced LTP was analysed by normalising the mean amplitudes from the last 10 minutes of baseline recordings after HFS (50-60 min post-HFS) to the mean of the amplitude of the initial 10 minutes baseline recordings before the HFS. The data from the control mice and the cKO mice were compared against pooled recordings from the independent control pathway that was generated by a second stimulation electrode located in the direction of the subiculum, as it cannot generate long-term plasticity changes in CA1. The Multi Channel Analyzer 2.2 (Multi Channel Systems MCS GmbH) software was used to record the peak value of each fEPSP amplitude. All data were analysed using the Multi Channel Analyzer 2.2 software, Microsoft Office Excel (Microsoft), and GraphPad Prism (GraphPad Software).

### Behavioral testing

Mice were handled for 2 min daily for 3-5 days before the start of the testing. The latter was performed between 8:00 A.M and 2:00 P.M, in sound-attenuating boxes under consistent lighting conditions (37 lux), except for the light-dark box test (600 lux). All mice were randomized and experimenters were blinded during the tests. The behavioral tests were videotaped (Basler camera) and mouse performance automatically tracked with EthoVisionXT software 15.0 (Noldus). Manual scoring of the recorded behavior was performed by a blinded experiment using The Observer XT12 software (Noldus Information Technology). Behavioral tests (see paragraph below). The tests were performed in the following order: OFT, NORT, three-chambers social interaction test, LD Box Test, and TWA test. Detailed descriptions of the protocols are provided in the supplementary information. Since social behavior was unaffected in the *Early postnatal deletion cohort* and in the *Adult deletion cohort*, the *Mouse line control cohort* was not assessed in the three-chambers social interaction test.

### Statistical analysis

The analyses were performed with GraphPad Prism software10.0 or with R software, version 4.4.2. Significance level was set to P < 0.05 and the data are presented as mean ± SEM. Sample distribution was assessed via Shapiro-Wilk normality test. For normally distributed dataset, two-group comparison was performed via unpaired or paired two-tailed t-test. For non-normally distributed dataset, two-group comparison was performed via Mann–Whitney U (unpaired data) or Wilcoxon test (paired data). Correlation was assessed via Pearson’s or Spearman’s (non-parametric) test. Comparison between more than two groups were carried out by, two-way or three-way ANOVA, followed by Tukey’s or Sidak post-hoc test. Mixed model design was applied for the statistical comparison of the nodes of Ranvier between groups. The number of biological replicates (n), specific statistical tests, and their P-values are indicated in the figure legends. Behavioral experiments were replicated in two independent batches. Detailed information of the statistical analyses is provided in the supplementary tables file.

## Acknowledgments

This work was supported by Leibniz Competition funds (Leibniz ScienceCampus NanoBrain, W71/2022), the German Research Foundation (CRC 1080, C02 to T.M.; PA 3913/2-1 to M.P.), Carl Zeiss Stiftung project InteReg (number P2024-02-015 to M.P. and A.W.) and Research Council of Finland (number 360360 to A. A.) The authors acknowledge Dr. R. Piccinno, Dr. S. Ritz and the IMB Microscopy & Histology Core Facility and Dr. Katrin Roth at the Core Facility Cellular Imaging, Center for Tumor Biology and Immunology of the Philipps University Marburg for supporting the image acquisition and analysis workflow. We are very grateful to Verena Opitz and Julia Deuster, for technical support during the experiments. We are grateful to Dr. Julia Leschik for helping to design the PCR primers. We are grateful to Dr. Konstantin Radyushkin, Dr. Ulrich Schmitt and the Mouse Behaviour Unit (MBU) of the Leibniz Institute for Resilience Research (LIR) for animal care and for supporting the animal experimental procedures. We are grateful to Prof. Jaqueline Trotter for the scientific discussion. We gratefully acknowledge Norarte Visual Science for the design of the graphical abstract. The work described herein was carried out in partial fulfilment of the requirements for a doctoral degree at the Johannes Gutenberg University Medical Center Mainz, Germany.

## Author Contributions

LM, MBM, GT designed research; LM, GP, CG, KB, LL, MP, JE, MK, DH, LN, AC, SW, CMG, LMT, HL, HS, JK, GT performed research; LM, GP, CG, KB, JE, AA, MP, SW, HS, GT analyzed data; LM, GP, CG, GT wrote the original draft of the paper; and LM, GP, CG, KB, LL, MP, JE, AA, MK, DH, LN, AC, SW, CMG, LMT, HL, HS, JK, RK, BL, AW, IH, TM, MS, MBM, GT reviewed and edited the original draft of the manuscript.

