## Supplementary file for "Glucocorticoid receptors in oligodendrocyte precursor cells regulate hippocampal network plasticity and stress-induced behavior in mice"

Univ.-Prof Dr. Giulia Treccani

Department of Systemic Neuroscience

Institute of Anatomy and Cell Biology

University of Marburg

Robert-Koch-Straße 8

35032 Marburg, Germany

**This PDF file includes:**

Supporting text
Figures S1 to S16
Tables S1
SI References

### Supplementary material and methods

#### NG2-CreER$^{T2}$ x Nr3c1$^{fl/fl}$ mouse line genotyping

Genotyping of NG2-CreER$^{T2}$ mice was performed using PCR with genomic DNA extracted from ear and tail samples. The following primers were used: 1 (5′ - GGC AAA CCC AGA GCC CTG CC - 3′), 2 (5′ - GCT GGA GCT GAC AGC GGG TG - 3′), and 3 (5′ - GCC CGG ACC GAC GAT GAA GC - 3′). These primers distinguish between the NG2 wild-type allele (557 bp) and the mutant allele (829 bp). Genotyping for Nr3c1$^{fl/fl}$ , to check for the presence of the loxP alleles (fl/fl) flanking the Nr3c1 gene (GR) and to confirm the deletion, the following primers were used: 1 (5′ - ATG CCT GCT AGG CAA ATG AT - 3′), 2 (5′- TTC CAG GGC TAT AGG AAG CA - 3′), and 3 (5′ -TGG TTG TTG GTG CTT TTG GCT AAA TC - 3′). These primers distinguish between homozygosity of the GR$^{wt/wt}$ alleles (250 bp), the GR$^{fl/fl}$ alleles (300 bp) and for the deletion (del) of the GR gene (GR fl/del), which shows an additional band at 400 bp (*SI Appendix*, Fig. S2 A).

#### Protein quantification via western blot

The samples were collected from mice that underwent behavioral testing. The entire cortex and hippocampus of 7 control (Ctlr) (4 males (M), 3 females (F)) and 7 cKO (4M, 3 F) were dissected, homogenized in ice-cold RIPA lysis buffer (ThermoFisher Scientific, 89900) and protease and phosphatase inhibitors (Roche, 04906837001, 04693132001), and centrifuged at 12,000 rpm for 20 min at 4 °C. Supernatants were collected and protein concentration assessed through bicinchoninic acid (BCA) assay (ThermoFisher Scientific, 23225). The samples were diluted in 4X Laemmli-buffer and incubated at 95°C for 5 minutes, except for the samples probed with oligodendroglial proteolipid protein (PLP) antibody, which were incubated with β-mercaptoethanol at 65°C. A total of 10-25 μg protein per sample (Fig.S6-S7) was separated on sodium dodecyl sulfate-polyacrylamide gel electrophoresis (SDS-PAGE, precast gels Bio-Rad, L007043). Proteins were then electrotransferred onto nitrocellulose membranes (GE Healthcare Life science, 10600002) in ice-cold transfer buffer and stained for total protein using Revert™ protocol (LiCor Biosciences). The membranes were blocked with 5% non-fat milk powder dissolved in Tris-buffered saline containing 0.05% Tween-20 (TBST) for 1 h at RT and subsequently incubated in primary antibody diluted in blocking solution at 4°C ON. The membranes were washed three times in TBST and then incubated with secondary antibody, diluted 1:10,000 in blocking solution, for 1 hour at RT. After three washes in TBST, signals were acquired using Odyssey Fc imaging system and image Studio 5.0 (LiCor Biosciences). Signal intensity for each sample was quantified by densitometric analysis and normalized to the corresponding total protein lane using Empiria Studio® Software 2.0 (LiCor Biosciences) (1). Data are expressed as a percentage of Ctrl. A list of the employed antibodies is provided in the supplementary material.

### List of employed antibodies

| AB | Company \| Provider | Cat. Nr. \| ID | Dilution |
|---|---|---|---|
| **Flow Cytometry** | | | |
| mouse anti-CD140a (Pdgfra) | Miltenyi Biotec | 130125521 | 1:10 |
| rat anti-CD45 | BioLegend | 103138 | 1:200 |
| rat anti-CD11b | BioLegend | 101215 | 1:1000 |
| mouse anti-GR | Santa Cruz Biotechnology | sc-393232 | 1:500 |

| | | | |
|---|---|---|---|
| chicken anti-CNPase | Sigma-Aldrich | AB9342 | 1:1000 |
| goat anti-mouse | Sigma-Aldrich | SAB4600066 | 1:500 |
| donkey anti-chicken | Sigma-Aldrich | SAB4600031 | 1:500 |
| **Histology** | | | |
| ***Primary*** | | | |
| rat anti-AN2 | Kind gift from Prof. J. Trotter, University of Mainz | | 1:250 |
| mouse anti-GR | Santa Cruz Biotechnology | sc-393232 | 1:250 |
| goat anti-Olig2 | RD Systems | AF2418 | 1:500 |
| rabbit anti-ASPA | Kind gift from Lutz Lab, University of Mainz | | 1:1000 |
| rabbit anti-NeuN | Abcam | ab177487 | 1:100 |
| rabbit anti-GFAP | Dako | Z0334 | 1:500 |
| rabbit anti-Iba1 | Wako | 019-19741 | 1:1000 |
| rabbit anti-PDGFRα | Cell Signaling Technology | 3174 | 1:200 |
| mouse anti-PDGFRβ | R&D Systems | AF1042 | 1:500 |
| rat anti-Ki-67 | ThermoFisher Scientific | SolA15. | 1:200 |
| mouse anti-CASPR | NeuroMab | #75-001 | 1:1000 |
| rabbit anti-Nav1.6 | Alomone labs | #ASC-009 | 1:250 |
| rat anti-Mbp | Serotec | MCA409S | 1:200 |
| ***Secondary*** | | | |
| goat anti-chicken | Aves Labs | F1005 | 1:500 |
| goat anti-rabbit | Invitrogen | A11011 | 1:500 |
| goat anti-mouse | Life Technologies | A11001 | 1:500 |
| goat anti-rabbit | Life Technologies | A11008 | 1:500 |
| goat anti-mouse | Invitrogen | A21236 | 1:500 |
| donkey anti-rabbit | Invitrogen | A31573 | 1:500 |
| donkey anti-goat | Life Technologies | A21432 | 1:500 |
| donkey anti-mouse | Life Technologies | A21202 | 1:500 |
| donkey anti-rabbit | Life Technologies | A21206 | 1:500 |
| goat anti-rat | Life Technologies | A21247 | 1:500 |
| donkey anti-mouse | Life Technologies | A31571 | 1:500 |
| goat anti-mouse | Abcam | ab150115 | 1:400 |
| goat anti-mouse | Life Technologies | A28175 | 1:400 |
| donkey anti-rat | Life Technologies | A21208 | 1:500 |
| **Western Blot** | | | |
| ***Primary*** | | | |
| rabbit anti-NG2 | Merck Millipore | AB5320 | 1:500 |

| | | | |
|---|---|---|---|
| rabbit anti-ASPA | Merck Millipore | ABN1698 | 1:1000 |
| mouse anti-CNPase | Merck Millipore | C5922 | 1:500 |
| rat anti-Myelin Basic Protein | Merck Millipore | MAB386 | 1:1000 |
| rat anti-PLP | Kind gift from Prof. E-M. Krämer-Albers(2) University of Mainz | cloneaa3 | 1:100 |
| ***Secondary*** | | | |
| goat anti-rat | LiCor Biosciences | 92568076 | 1:10000 |
| goat anti-rat | LiCor Biosciences | 92532219 | 1:10000 |
| goat anti-rabbit | LiCor Biosciences | 92532211 | 1:10000 |
| goat anti-mouse | LiCor Biosciences | 92532210 | 1:10000 |

### Microscopy and image analysis

### Myelin content and internode length

For the analysis of myelin content and internode length, confocal micrographs were acquired with CS2 63x/1.40 oil objective, at a format of 1024 × 1024 pixels, the pixel size to 101nm x 101nm and the z-stack interval to 150nm. The pinhole was set to 1 AU. The size of the overall field of view, acquired via tail scan, was 288.50um x 195.33um. For each mouse, 2 micrographs (1 brain section, both hemispheres) from the dorsal CA1 and L2/3 of the somatosensory cortex, were acquired and Z-stack images were processed and analyzed with ImageJ (NIH) software by a blinded experimenter (2) and with IMARIS Software (https://imaris.oxinst.com). For the overall myelin content quantification, the optical plane with the maximum mean grey value was identified. This optical plane and the 10 optical planes above and below were included in the analysis, so that 6 µm of the total section thickness was used for quantification. Semi-automated segmentation of Mbp+ fibers was performed via a custom-written macro and the area covered by the MBP+ signal was employed as proxy for myelin content (3).

For the analysis of the internode length, semi-automated 3D surface reconstruction of Mbp signal was performed in IMARIS. Internodes flanked by paranodal structures at both extremities were manually identified and labelled by a blinded experimenter. The length of this subset of paranode-flanked internodes was estimated via *BoundingBoxOO length C* parameter.

**Analysis of paranodes and nodes of Ranvier**

For the analysis of paranodes and nodes of Ranvier, imaging was performed with ZEISS LSM 880 Confocal Laser Scanning microscope with AiryScan (Zeiss). To image individual nodes within a field of view, a region around a node was cropped, and a z-stack of the cropped region was acquired. Z-stacks were acquired and pixel size set at 0.04 × 0.04 × 0.10 μm (x,y,z). For each mouse, 4-5 brain sections including CA1 region were analysed. An automated pipeline to segment and analyze the length of paranodes and nodes of Ranvier in the acquired 3D microscopy images was implemented (4).

**OPC morphology: confocal acquisition and analysis parameters**

For the confocal acquisition, the field of view was located so that the soma of the NG2-cell of interest was in the center of the micrograph. A minimum of ten cells for each mouse was acquired by a blinded experimenter. The semi-automated segmentation of the cell processes was performed with IMARIS software 10.2.0 (https://imaris.oxinst.com) through the Filament Tracing Wizard (Detection type: Starting Point; Starting point diameter: 10um) combined with AI-training based-Seed Point Classification (Segment Seed Point diameter: 0.5um). The reconstruction accuracy was evaluated by a blinded experimenter and manual refinement of the segmentation was carried out in the rare cases that required it. A final filtering based on the *filament length sum* was employed to exclude from the analysis any artefact and/or process that did not belong to the analyzed cell. The employed algorithm settings are outlined below.

[Algorithm]
Name = Autopath (no loops)
Segment Start Point = true
Detect Spines = false
Enable Regions of Interest = false
Track (over time) = false
[Soma Starting Points]
Segment Channel Index = 2
Segment Starting Point Diameter = 10.0 µm
Calculate Soma Model = true
[Filter Starting Points]
Segment Starting Point Threshold Low = 9.321
Segment Starting Point Threshold High = Automatic
Render Soma Model = false
[Seed Points for Segments]
Segment Seed Point Diameter = 0.500 µm
[Filter Seed Points for Segments]
Segment Seed Point Threshold = 33.886
Diameter around Starting Point(s) to remove Seed Points = 15.0 µm
Segment Diameter Filter Strengthness = 2
[Seed Points Classification]
Group Name = Filter
Input = All Spot
No. of Classes = 2
Class:: Name = Keep
Color = 0.000 1.000 1.000
Class:: Name = Discard
Color = 1.000 0.000 0.000
FilterType = ML
Training Data
Class:: Name = Keep
Size = 22
Class:: Name = Discard
Size = 48
[Segment Classification]
Group Name = Filter
Input = All Segment
No. of Classes = 2
Class:: Name = Keep
Color = 0.000 1.000 1.000
Class:: Name = Discard
Color = 1.000 0.500 0.000
FilterType = ML

Training Data
Class:: Name = Keep
Size = 93
Class:: Name = Discard
Size = 63
[Tree Build]
[Terminal Segment Postfilter]

**Electrophysiological Recordings – Multielectrode Array**

Naïve mice were employed for this experiment. Three adult mice per group and per sex were employed. For each mouse, 3-9 paired recordings were taken, with each recording coming from a new individual slice. The mice were deeply anesthetized (i.p. 200 mg/kg ketamine) and transcardially perfused with choline-based artificial cerebrospinal fluid (cACSF) solution [125 mM NaCl, 3 mM KCl, 1.3 mM, $MgSO_4$ x $7H_2O$, 2.5 mM, $CaCl_2$ x $H_2O$, 26 mM $NaHCO_3$, 37.5 mM choline chloride and 13 mM d-glucose, pH of 7.4]. The mouse was then decapitated, and the brain was quickly removed and placed into icy (4°C) cACSF perfused with carbogen (95% oxygen ($O_2$) and 5% carbon dioxide ($CO_2$)). After 1 minute, 350µm-thick sections were collected and placed in normal ACSF at RT for at least 40 minutes before use. Individual slices were then transferred to a multielectrode array (MEA) chip of a two-chamber MEA system (MEA2100 System, Multi-Channel Systems MCS GmbH; Chip: 60MEA200/30iR). Here, the slice was constantly perfused with carbogenated normal ACSF [125 mM NaCl, 3 mM KCl, 1.3 mM $MgSO_4$ x $7H_2O$, 2.5 mM $CaCl_2$ x $H_2O$, 1.25 mM $NaH_2PO_4$ x $H_2O$, 26 mM $NaHCO_3$ and 13 mM d-glucose at a temperature of 32°C. The normal ACSF also contained 0.01% EtOH to control for any ethanol-induced electrophysiological changes (see below). The stimulating electrode was positioned on CA3 region and the recording electrode in the CA1 region of the hippocampus. The recording was performed along the Schaffer Collateral pathway. All electrophysiological recording protocols were generated and applied by Multi Channel Experimenter 2.2 software (Multi Channel Systems MCS GmbH), which had a 50 kHz sampling rate and a 200Hz cut-off Butterworth highpass 2.0 Order Filter. The recording of the reported outputs was performed and quantified via Multi-Channel Analyzer 2.2 (Multi Channel Systems MCS GmbH). Spontaneous neuronal activity was identified as an event (recorded in 5-min traces) during which the extracellular spike frequency exceeded the downward threshold of five standard deviations from baseline activity noise. For evoked activity, an input-output curve was generated by applying stepwise voltage stimulations (0.5V to 5V in 40 second-separated 500mV steps). The maximum amplitude of each evoked field excitatory postsynaptic potential (fEPSP) was recorded for each individual slice. For the paired-pulse stimulation, a voltage stimulation intensity to produce a fEPSP equal to approximately 30% of the maximum evoked fEPSP amplitude (see “input/output curve” above) was applied. A 50ms inter-stimulus interval was employed. From there, the amplitude of the second response was divided by the first to obtain the paired-pulse ratio. After these control recordings, the perfusion input was switched to carbogenated ACSF containing 200nM of Dexamethasone diluted in ethanol (0.01%). After a wash-in of 30 minutes, all protocols were repeated in the same fashion as described above on the same slice. On finishing, the slice was removed and the drug-infused ACSF was completely washed out, before placing the next slice. All electrophysiological data were analyzed using the Multi-Channel Analyzer

2.2 software, Microsoft Office Excel (Microsoft), and Prism software (GraphPad 10.0).

**Behavioral testing – detailed**

*Open field test (OFT).* The mice were placed in the center of a square, non-transparent polyvinyl chloride (PVC) arena (44 cm x 44 cm x 40 cm) and allowed to explore freely for 10 minutes. Total distance travelled was scored.

*Novel object recognition test (NORT).* In the same OFT arena, the experimental mouse was allowed for 10-min exploration of two identical objects placed at equidistant locations (*training phase*) and was then returned to the homecage. After 24-hour, the mouse was reintroduced into the arena for 5-min exploration of one familiar and one novel object, which were similar in size but differed in shape, material, texture, and contrast (*testing phase*). Object exploration was defined as direct or proximal interactions with the object, including sniffing or touching it with the nose or forepaws. The ratio of time spent exploring the novel object to the total exploration time of both objects was calculated as the novel object recognition index (expressed as a percentage). Trials in which the total exploration time was less than 5 seconds were considered insufficient and excluded from the analysis.

*Three-chambers social interaction test.* The three-chambers social interaction test was conducted as previously described (5). Briefly, the experimental mouse was habituated for 5 minutes to the central chamber of a three-chambered apparatus.

Afterwards, while the experimental mouse was briefly returned to the homecage, a same-sex juvenile (or young adult in case of the *adult deletion cohort*) C57BL/6 mouse (social target) was placed under one mesh cylinder located in one of the two outer chambers of the apparatus and a metal ring (non-social target) under a meshed cylinder in the other outer chamber. The test mouse was returned to the three-chambered apparatus and allowed to freely explore all three chambers for 5 minutes. The activity was videotaped and tracked with a dedicated video-tracking software. Social interaction, defined as any nose entry into a pre-defined interaction zone around each cylinder, was scored by a blinded experimenter.

*Light-Dark (LD) Box Test.* The LD box test was carried out as previously described (6). Briefly, mice were placed in the lit compartment facing the wall and observed for 5 minutes. The percentage of time spent in the light, the latency to enter the dark compartment, and the frequency of light-dark transitions were analyzed.

*Two-way active avoidance (TWA) test.* The TWA test was performed using a multi-conditioning system (TSE systems) as previously described (7, 8). Briefly, the mice received one training session per day for five consecutive days. Each session consisted of 60-sec habituation followed by 25 learning trials. Each trial included a 5-sec 1 kHz 60 dB tone (conditioned stimulus, CS) immediately followed by a 0.4 mA E-shock (unconditioned stimulus, US; cut-off: 5-sec), and an additional 60-second delay. The E-shock ceased either when the mouse moved to the other compartment or after 5-sec, if no movement occurred. If the mouse moved during

CS the E-shock was not delivered. An inter-trial interval of 30 seconds was maintained between trials. A conditioned response (CR) implicated an active transfer to the other chamber upon CS presentation; an unconditioned response (UR) implicated an active transfer in response to the US; a failure consisted of the absence of any transfer to the alternate compartment. Learning performance was calculated as the percentage of CR relative to total responses (CR and UR) for each animal in each trial. Mice repeatedly escaping from the test box during trials were excluded (n=2).

**Automated behavioral screening via DeepLabCut**

The automated behavioral screening was performed with DeepLabCut (9). Each original video (1280 x 1024 pixels, 25 f/s) was cropped and cut to a final format of 256 x 256 pixel and 5 min duration. DeepLabCut was trained to detect and discriminate between the snout, left ear, right ear, and tail base of the mouse in the video. The data were filtered for likelihoods greater than 0.95. and the center point was calculated and exported. The output data included the following items: batch, trial, and arena; for each trial time and xy coordinates were extracted and ID, genotype, sex, and a "switched" attribute were added. The switched attribute identifies the videos where the left toy was replaced with a new one in the testing of the NORT. The traces (i.e., position of the mouse over time) were visualized as line plot and heatmap and were overlaid on the video. The Euclidean distances to the objects were calculated and added to the table for each position (dl/r: position to left/right object [px]). The speed for consecutive positions was calculated as pixels/minute; the absolute angle between the movement and the direction to each object was calculated (al/r: angle to left/right toy [degrees]).

The angle values of each mouse position (range: 0-180°) were normalized to values from 1 to 0, with 1 indicating direct approach and interaction with the object, and 0 indicating retreat from the object (al/rn: angle left/right normalized). The distance to the object values were normalized so that they would range from 1 to 0, with 1 indicating a direct approach to the object, and 0 representing the maximum distance from the object (dl/rn: distance left/right normalized). The normalized distances and angles were categorized into equal bins for frequency counting (dn/otn[0-9]: distance new/old object normalized bin #, angn/otn[0-9]: angle new/old object normalized bin #). The difference in frequency between the bins for the new and old object was analyzed for angles and distances (d/ant[0-9] – d/aot[0-9]: distance/angle new toy bin – distance/angle old toy).

**Supplementary figures**

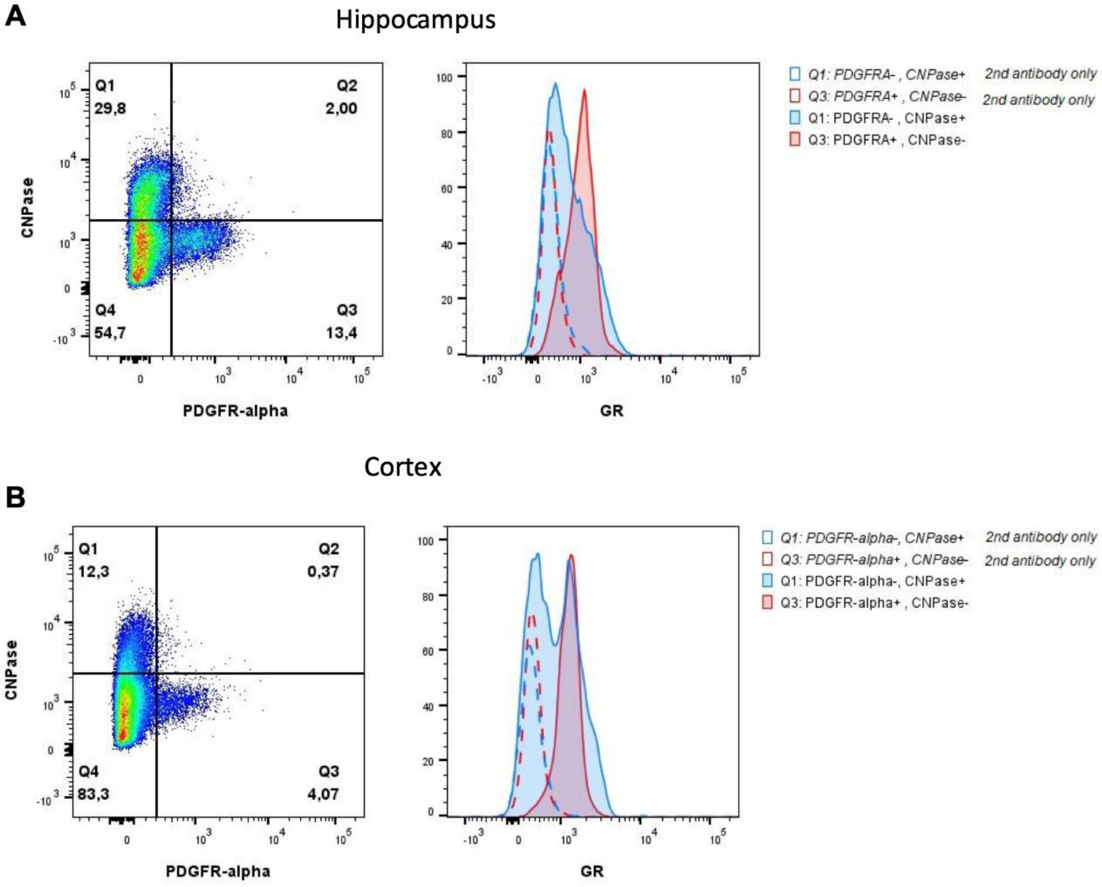


**Fig. S1. Glucocorticoid receptor (GR) expression by mature oligodendrocytes and their precursors in the cortex and hippocampus of adult mice.** Single-cell suspensions from dissected hippocampi (**A**) and cortex (**B**) were analyzed by flow cytometry. After excluding dead cells, doublets, and CD45+CD11b+ cells, mature oligodendrocytes and precursor cells were identified by expression of CNPase and PDGFR-alpha, respectively (dot-plots, left panel). Each population was then displayed in overlayed histogram plots indicating GR expression (right panel): CNPase+ (blue) and PDGFRA+ (red) populations with (filled) or without (secondary antibody only, dashed) anti-GR antibody.

Figure S2

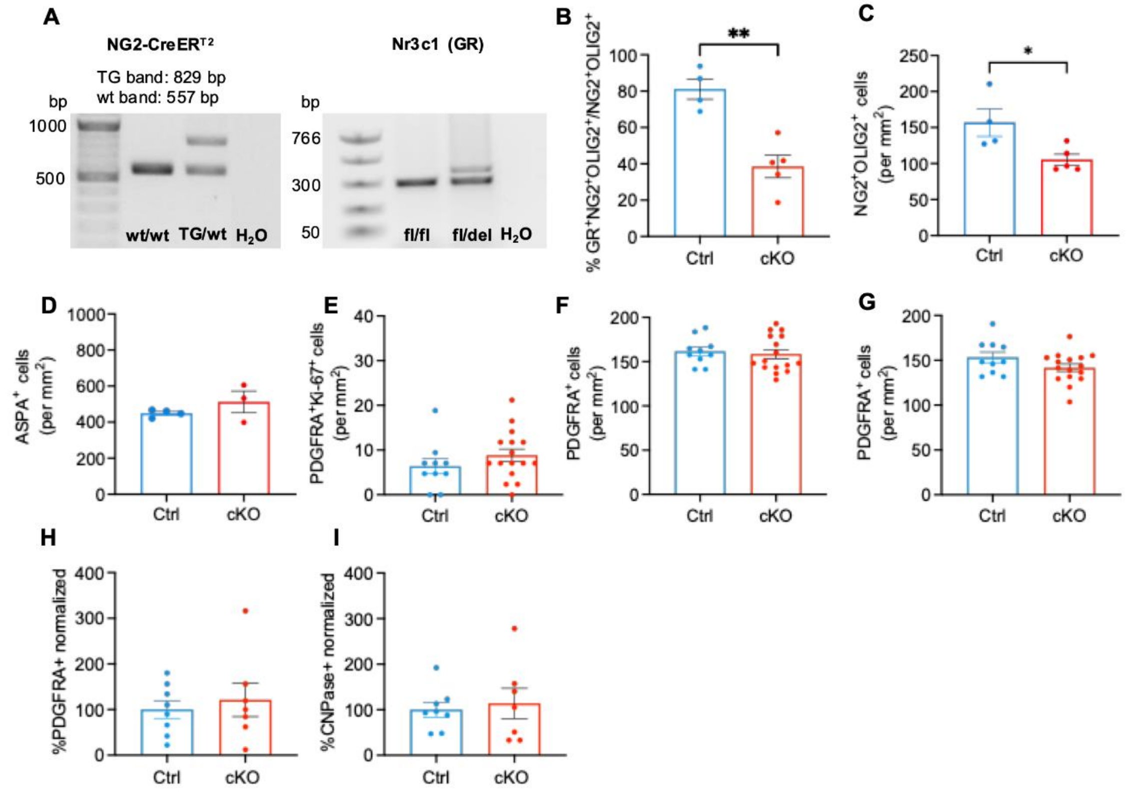


**Fig.S2. Generation of NG2-CreERT2; Nr3c1fl/fl mice and quantification of OPCs and oligodendrocyte density in the cortex of adult mice** (**A**) PCR genotyping of knock-in mice expressing Cre DNA recombinase variant CreERT2 in the NG2 (Cspg4) locus, and the floxed Nr3c1 (GR) gene. (**B-G**) Quantification of Cre recombinase efficiency and OPCs and myelin marker density in Ctrl and cKO mice. (**B**) Percentage of GR+NG2+ cells the cortex (unpaired t test: $t(7) = 4.956$, $p = 0.0016$; n = 4 Ctrl and n = 5 cKO mice). (**C**) Density of NG2+Olig2+ cells (unpaired t test: $t(7) = 2.718$, $p = 0.0299$; n = 4 Ctrl and n = 5 cKO mice). (**D**) Density of ASPA+ cells in the cortex (unpaired t test with Welch's correction: $p = 0.4041$; n = 4 Ctrl and n = 3 cKO mice). (**E**) Density of PDGFRA+Ki-67+ cells in the cortex (unpaired t test: p = 0.2725; n = 10 Ctrl and n = 16 cKO mice). (**F**) Density of PDGFRA + cells in the hippocampus (unpaired t test: p = 0.683), and **(G)** in the cortex (unpaired t test: p = 0.1626; n = 10 Ctrl and n = 16 cKO mice). (**H**) Percentage of single PDGFRA + cell population among CD45-CD11b- cells dissected from the cortex (unpaired t test: p = 0.6042). (**I**) Percentage of single CNPase+ cell population among CD45-CD11b- cells dissected from the cortex (unpaired t test: p = 0.7033; n = 8 Ctrl and n = 7 cKO mice). Data are expressed as the mean ± S.E.M. and each dot represents one animal. *P < 0.05, **P <, 0.01.

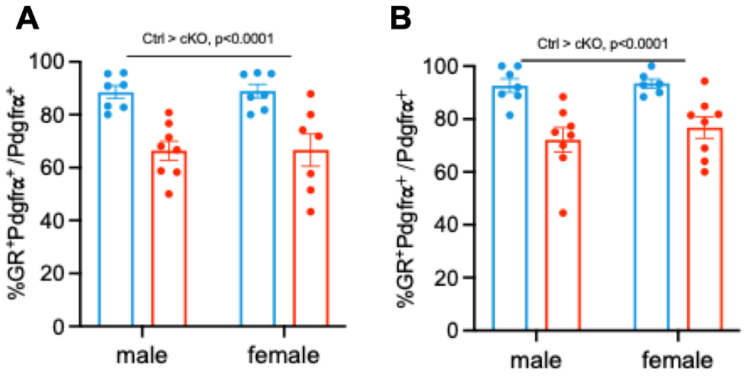


**Fig. S3. The loss of GR in OPCs and in mature oligodendrocytes is equally observed in both sexes.** (**A**) Percentage of GR+-OPCs with respect to the overall OPC population in the hippocampus (two-way ANOVA, genotype: F (1, 25) = 31.75, p<0.0001; sex: F (1, 25) = 0.008369, p =0.9278; genotype x sex: F (1, 25) = 0.008369, p=0.9888). (**B**) Percentage of GR+-OPCs with respect to the overall OPC population in the cortex (two-way ANOVA, genotype: F (1, 25) = 24.58, p<0.0001; sex: F (1, 25) = 0.5317, p =0.4727; genotype x sex: F (1, 25) = 0.2711, p=0.6072). Data are expressed as the mean ± S.E.M. and each dot represents one animal.

Figure S4

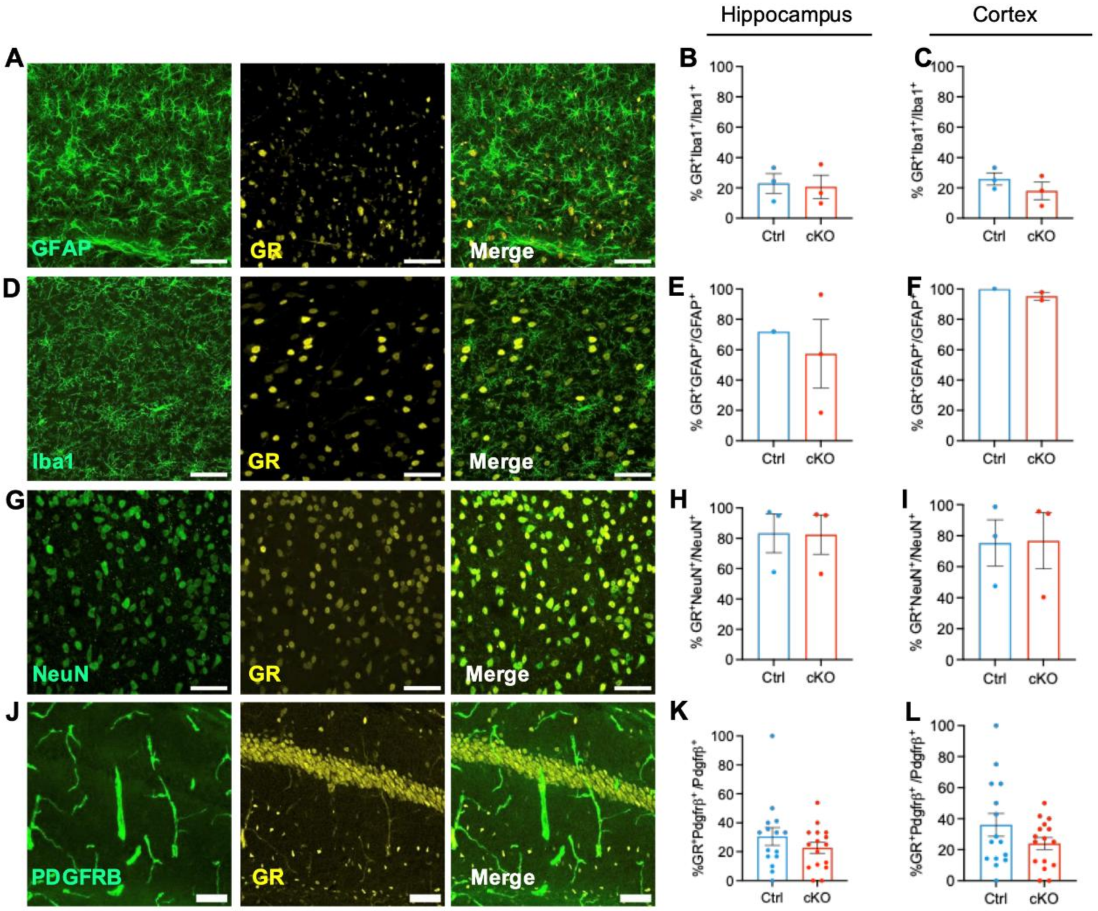


**Fig.S4. GR expression in astrocytes, microglia, neurons and pericytes in cKO mice**. (**A-C**) Representative confocal images and quantification of the co-expression of GR with astrocytes (GFAP) in hippocampus and cortex, n = 1 Ctrl and n = 3 cKO. **(D-F)** Representative confocal images and quantification of the co-expression of GR with microglia (Iba1) in hippocampus and cortex, n = 3 Ctrl and n = 3 cKO. **(G-I)** Representative confocal images and quantification of the co-expression of GR with neurons (NeuN) in hippocampus and cortex, n = 3 Ctrl and n = 3 cKO. **(J-l)** Representative confocal images and quantification of the co-expression of GR with pericytes (PDGFRB) in hippocampus (Mann-Whitney,

U=95.50, p=0.3407) and cortex (Unpaired t test with Welch's correction, t=1.454, df=21.10, p=0.1606), n = 15 Ctrl and n = 16 cKO. Scale bars = 50⌠m. Images acquired in Cornu ammonis 1(CA1) and layer 2/3 of the primary somatosensory cortex, Bregma: -1.79 to -2.53 mm. Data are expressed as the mean ± S.E.M and each dot represents one animal. Brightness and contrast of the micrographs have been adjusted for display purposes.

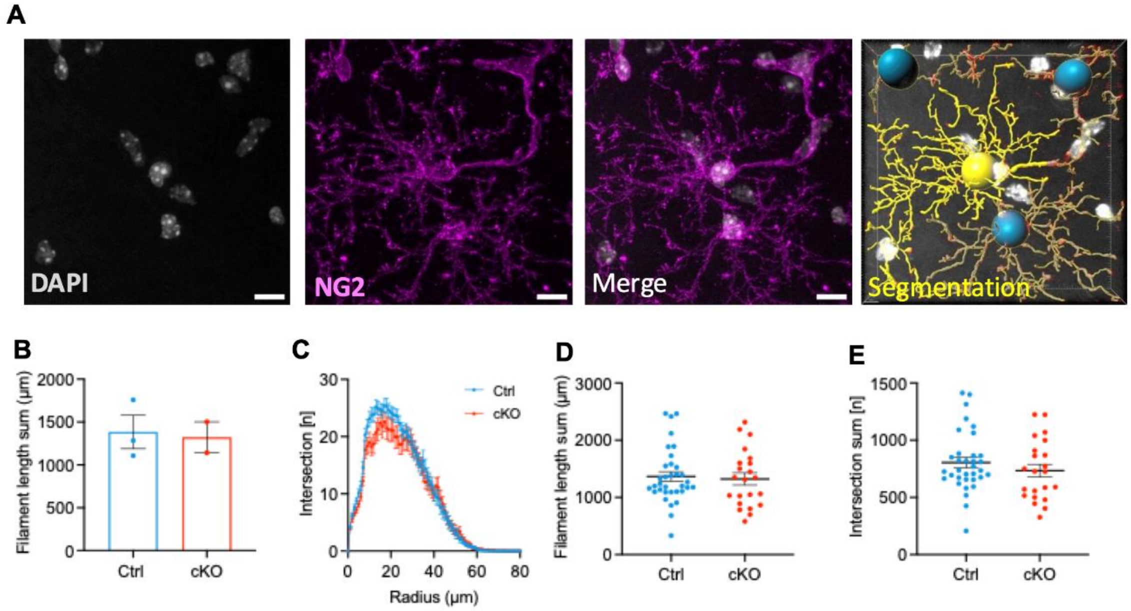


**Fig. S5. Morphological characterization of OPCs in cKO mice.** (**A**) Representative images of NG2+ cell with segmentation analysis micrograph in hippocampal CA1 subregion. Scale bar = 10 µm. Brightness and contrast have been adjusted for display purposes. (**B**) Mean of filament length sum and (**C**) OPCs branch intersections. (**D**) Filament length sum distribution. (**E**) Intersection sum distribution. Sample size: Ctrl, n=3 (cell, n=34); cKO, n=2 (cell, n=23).

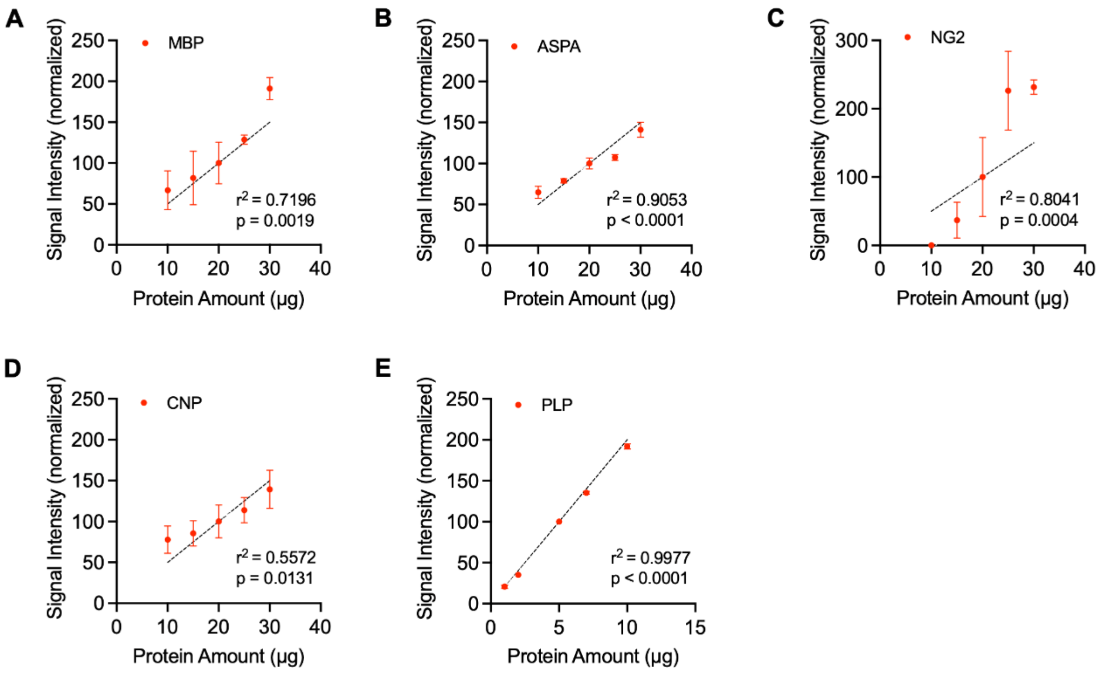


**Fig. S6. Evaluation of the linearity in protein-level quantification.** The correlation between the amount of loaded protein and the signal intensity is linear within the range utilized to compare Ctrl and cKO mice. Samples from two Ctrl mice. For MBP (**A**), ASPA (**B**), NG2 (**C**), and CNP (**D**), 10, 15, 20, 25, and 30 μg of total protein were loaded. For PLP (**E**), 1, 2, 5, 7, and 10 μg of total protein were loaded. Data points represent the average intensity of all two samples at each amount of total protein (normalized to the signal intensity at 20 μg or 5 μg total protein). The intensity for NG2 deviated slightly from the linear relation, but both signals still showed significant differences with 25% changes in total protein levels. Simple linear regression, error bars represent S.E.M. The dashed line represents the linear regression plot, predicting signal intensities of 50% at 10 μg, 75% at 15 μg, 100% at 20 μg, 125% at 25 μg, and 150% at 30 μg. The measured signal intensity for the synaptic proteins of interest closely aligned with this hypothetical plot within the range of 20 to 25 µg or 2,5 to 10 µg total protein which was used for the study, confirming the linearity of the protein quantifications (**A-E**). PLP total protein stain for normalization purposes showed a tight linear relationship with the amount of total protein loaded (**E**).

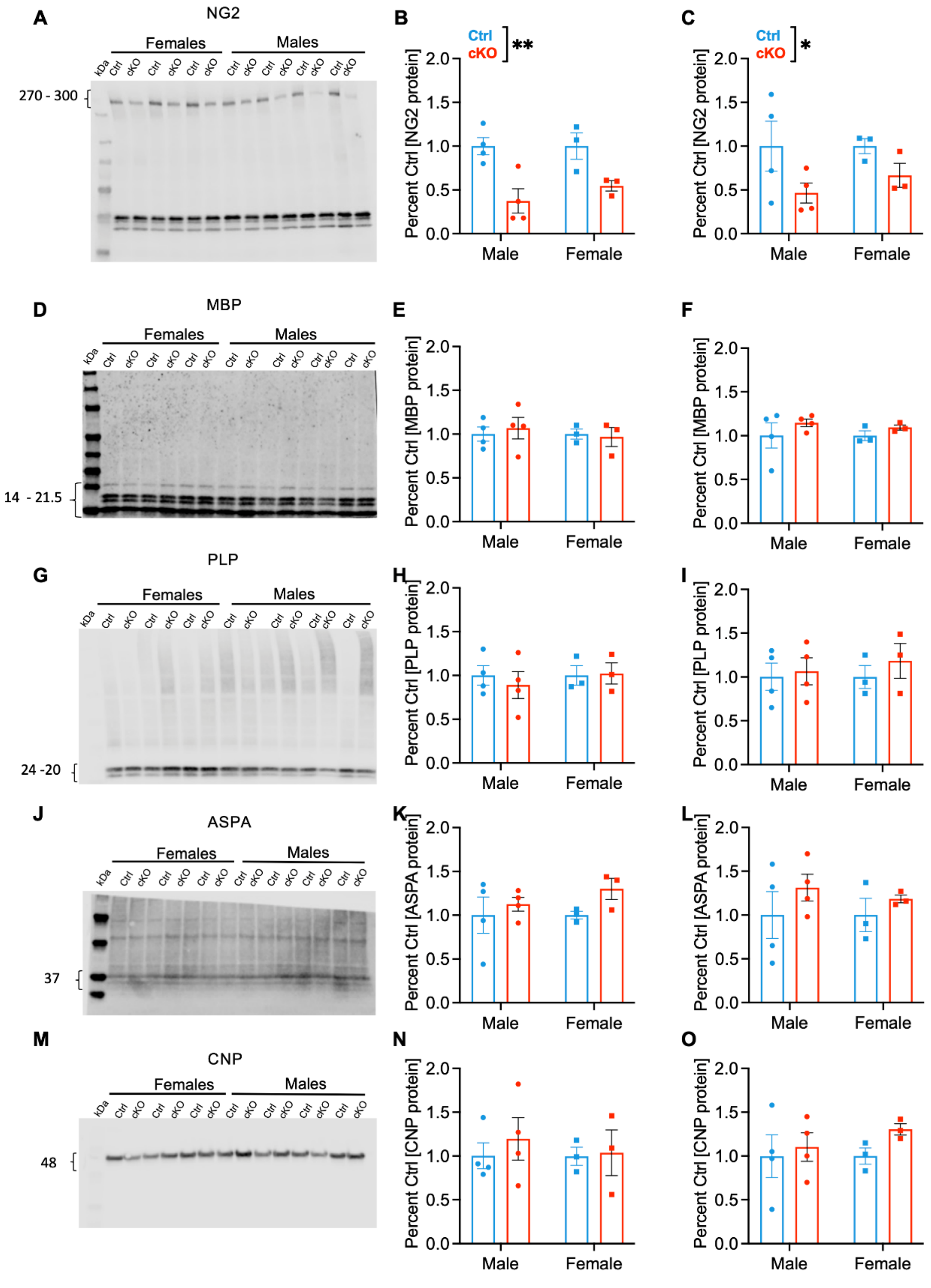


**Fig. S7. Quantification of OPCs and myelin markers proteins level in the hippocampus and cortex.** Representative western blotting of Hippocampi (HP)

or cortices (CX) lysates stained for oligodendrocyte lineage-related markers (n = 4M/3F Ctrl and n= 4M/3F cKO mice). (**A-C**) **Neural/glial antigen 2 (NG2)** (**A**) Representative blot. (**B**) NG2 protein level in HP (Two-way ANOVA: genotype, F (1, 10) = 19.76, p = 0.0012; sex and interaction, p = 0.4953). (**C**) NG2 protein level in CX (Two-way ANOVA: genotype, F (1,10) = 5.060, p = 0.0482; sex and interaction, p ε 0.6128). (**D-F**) **Myelin basic** protein**.** (**D**) Representative blot. (**E**) MBP protein level in HP (Two-way ANOVA: p = 0.6299. (**F**) MBP protein level in CX (Two-way ANOVA: p = 0.2279. (**G-I**) **Proteolipid protein (PLP)**. (**G**) Representative blot. (**H**) PLP protein level in HP (Two-way ANOVA: p = 0.623). (**I**) PLP protein level in CX (Two-way ANOVA: pε0.4687). (**J-L**) **Aspartoacylase (ASPA)**. (**J**) Representative blot. (**K**) ASPA protein level in HP (Two-way ANOVA: p ε 0.1643). (**L**) ASPA protein level in CX (Two-way ANOVA: p = 0.2442). (**M-O**) **2′,3′-cyclic nucleotide 3′-phosphodiesterase, E.C.3.1.4. 37 (CNP).** (**M**) Representative blot. (**N**) CNP protein level in HP (Two-way ANOVA: p = 0.5839. (**O**) CNP protein level in CX (Two-way ANOVA: p = 0.2822). Data are expressed as the mean ± S.E.M and each dot represents one animal. *P < 0.05, **P < 0.01.

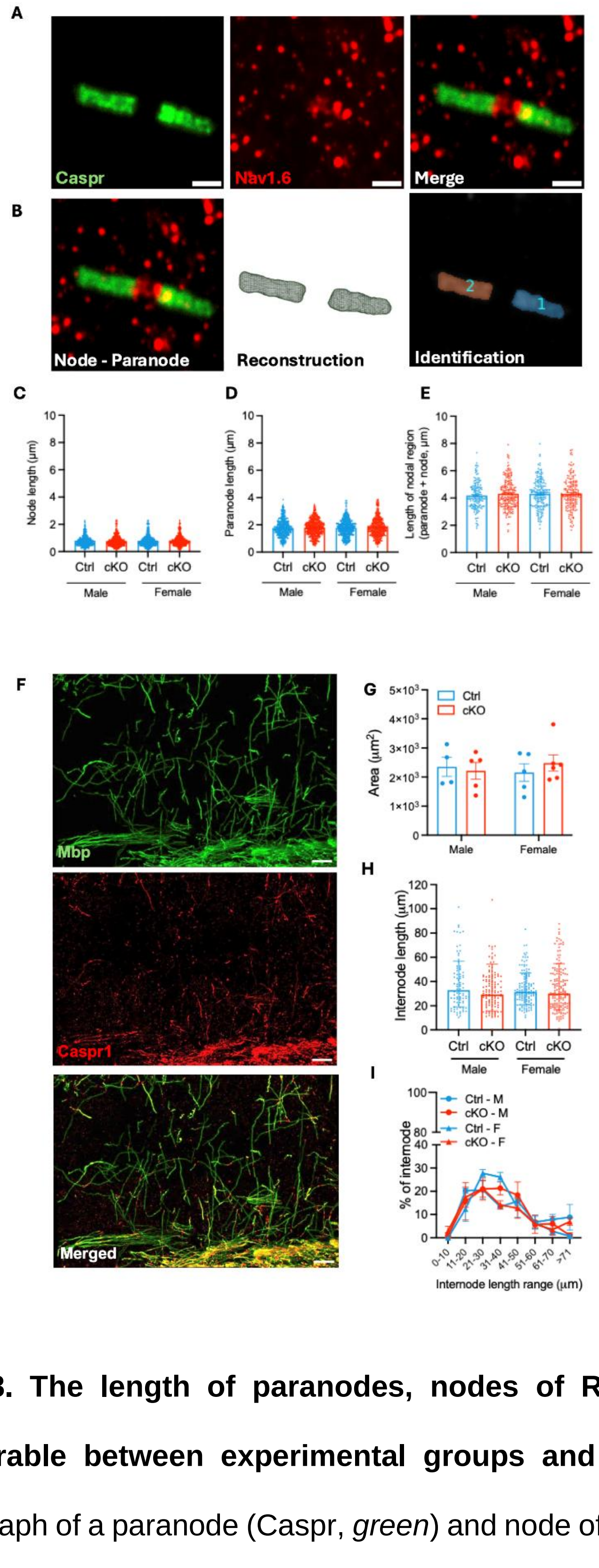


**Fig. S8. The length of paranodes, nodes of Ranvier, and internodes is comparable between experimental groups and sexes.** (**A**) Representative micrograph of a paranode (Caspr, *green*) and node of Ranvier (Nav1.6, *red*); scale bar = 1 μm. Brightness and contrast have been adjusted for display purposes. (**B**) Representative images of the reconstruction and identification of two paranodes and the related node of Ranvier. (**C**) Node length (Linear mixed model, p>0.05; Ctrl male, n = 390 nodes; cKO male, n = 481 nodes; Ctrl female, n = 465 nodes; cKO female, n = 456 nodes). (**D**) Paranode length (Linear mixed model, p>0.05; Ctrl male, n = 444 nodes; cKO male, n = 584 nodes; Ctrl female, n = 515 nodes; cKO female, n = 517 nodes). (**E**) Length of paranodes and the related node of Ranvier (Linear mixed model, p>0.05; Ctrl male, n = 157 nodes; cKO male, n = 187 nodes; Ctrl female, n = 209 nodes; cKO female, n = 174 nodes). Graph shows mean ± S.E.M, total mice, n= 28 (Ctrl male, n = 6; cKO male, n = 8; ctrl female, n = 7; cKO female, n = 7). (**F**) Representative confocal images, scale bar = 20 μm. (**G**) Myelin content in hippocampal CA1 (Two-Way ANOVA: Sex F (1,16) = 0.01383, p = 0.9078; genotype, F (1, 16) = 0.09833), p = 0.7579; sex x genotype, F (1, 16) = 0.5999, p = 0.4499). (**H**) Internodal length in hippocampal CA1 (Two-Way ANOVA: Sex F (1, 16) = 0.04783, p = 0.8296; genotype, F (1, 16) = 0.5837, p = 0.4560; sex x genotype, F (1, 16) = 0.2305, p = 0.6377). (**I**) Binned internodal length in hippocampal CA1 (Three-way ANOVA: Internode range, F (7, 112) = 19.58, p<0.0001; sex, F (1, 16) = 2.898, P=0.1080; genotype, F (1, 16) = 1.619, P=0.2214; Internode range x sex, F (7, 112) = 0.6832, P=0.6860; Internode range x genotype, F (7, 112) = 0.2093, P = 0.9827; sex x genotype, F (1, 16) = 0.06740, P=0.7985; Internode range x sex x genotype, F (7, 112) = 2.086, P=0.0507. Data are expressed as the mean ± S.E.M, except for (**H**), where data are expressed as the geometric mean ± S.E.M, *P<0.05, **P < 0.01. Contrast and brightness of the confocal images have been adjusted for display purposes. *Males:* n = 4 Ctrl and n = 5 cKO; *Females*: n = 5 Ctrl and n = 6 cKO.

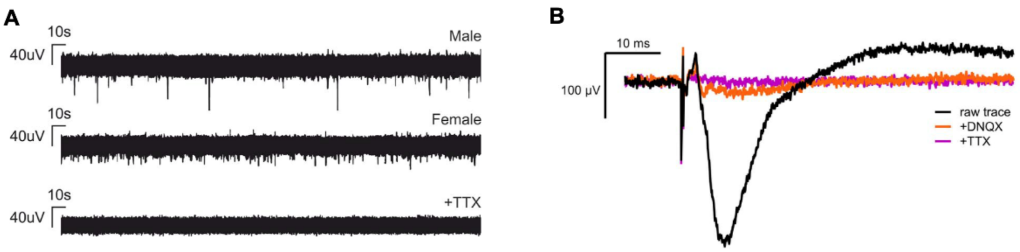


**Fig. S9. DNQX/TTX control experimental traces.** (**A**) Representative five-minute traces from spontaneous spiking activity recordings of a male and female mouse, and a trace showing the elimination of spiking activity following a 10-minute wash-in of tetrodotoxin (TTX). (**B**) Overlayed evoked fEPSP traces showing the raw signal (black), after 10-minute wash-in of 6,7-dinitroquinoxaline-2,3-dione (DNQX) (orange) and an additional 10-minute wash-in of TTX (pink).

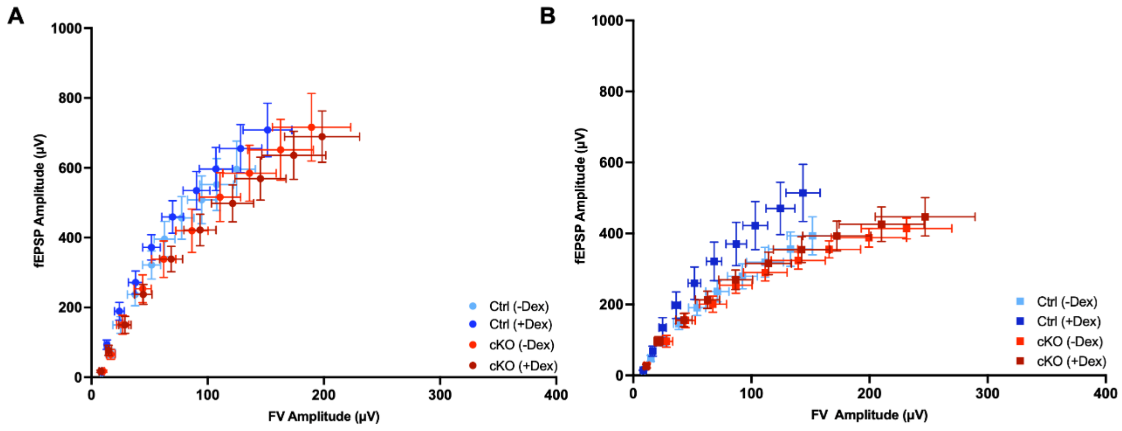


**Fig. S10**. **Fiber volley**. (**A**) Fiber Volley (FV) Amplitude vs. fEPSP Amplitude in male Ctrl vs. cKO with and without dexamethasone treatment. (**B**) FV Amplitude vs. fEPSP Amplitude in female Ctrl vs. cKO with and without dexamethasone treatment. Graph shows mean ± S.E.M.

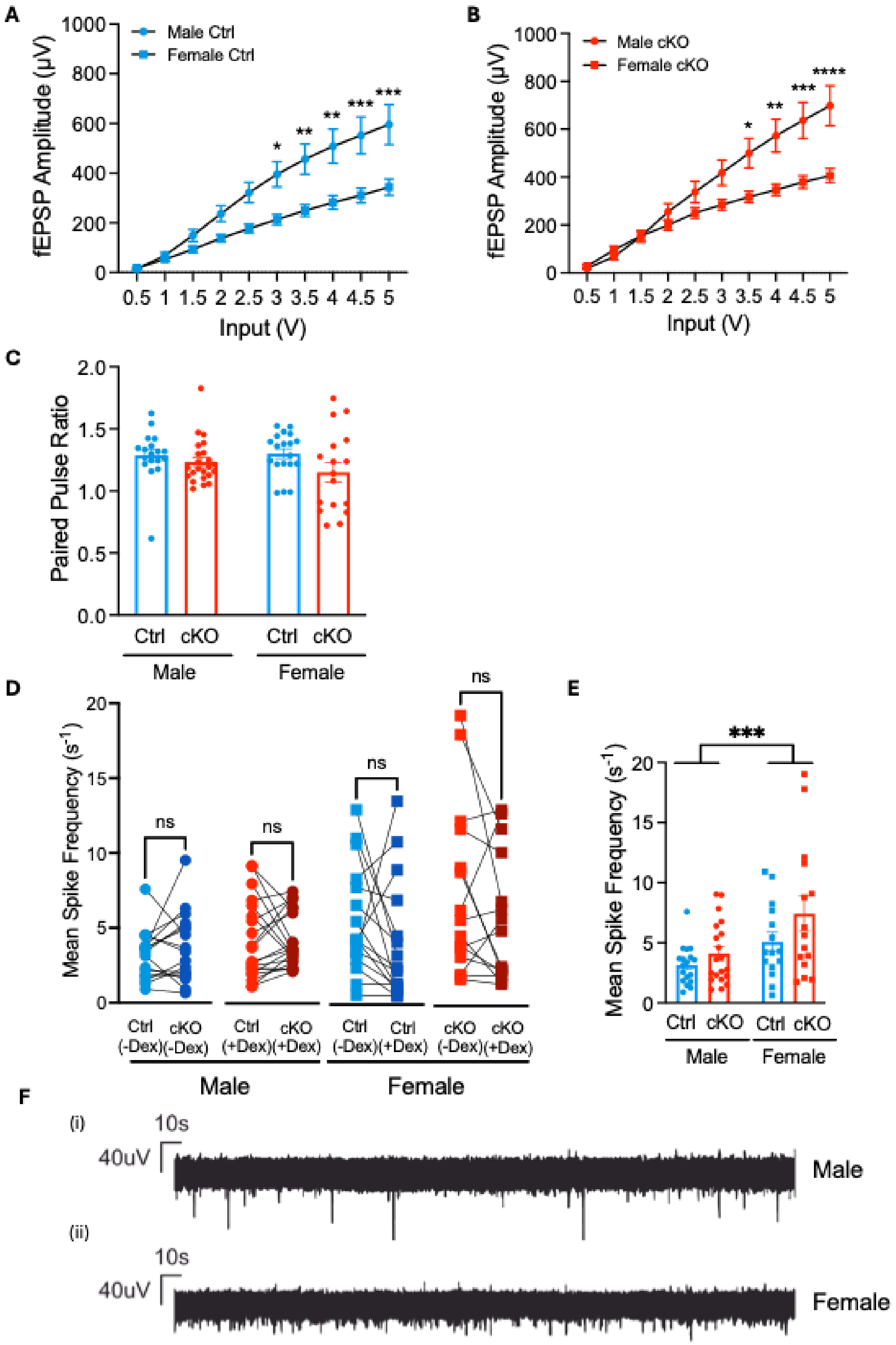


**Fig. S11. Male mice show higher hippocampal network excitability than female mice under resting conditions and acute stress failed to alter**

**spontaneous network activity of either sex or genotype in the hippocampal CA1 region, but a sex-specific difference was found at the basal status before drug application**. (**A**) Evoked field potential in the CA1 regions of males vs. females Ctrl mice. (**B**) Evoked field potential in the CA1 regions of males vs. females cKO mice. (***Ctrl***: Two-Way RM ANOVA: Input x sex, $F(9, 288) = 7.622$, $p < 0.0001$; Šídák's multiple comparisons test: Input 3: M > F, $p = 0.0185$; Input 3.5: M > F, $p = 0.0043$; Input 4: M > F, $p = 0.0012$; Input 4.5: M > F, $p = 0.0004$; Input 5: M > F, $p = 0.0002$; n = 18 male and n = 16 female. ***cKO***: Two-Way RM ANOVA: Input x sex, $F(9, 333) = 9.396$, $p < 0.0001$; Šídák's multiple comparisons test: Input 3.5: M > F, $p = 0.0412$; Input 4: M > F, $p = 0.0039$; Input 4.5: M > F, $p = 0.0006$; Input 5: M > F, $p < 0.0001$; n = 22 male and n = 17 female). (**C**) Baseline paired pulse ratio (Two-way ANOVA: genotype, $F(1, 73) = 3.934$, $p = 0.0511$; sex and interaction, p ε 0.3529). (**D**) Paired recordings within sex and genotype of the mean spike frequency of the hippocampal region before and after the 30-minute wash-in of dexamethasone (Wilcoxon matched-pairs signed rank test or paired t test: pε 0.1297). (**E**) Spontaneous spiking at basal state. (**F**) Representative trace of a 5-minute recording showing male (i) and female (ii) spiking activity without any drug wash-in. Graph shows mean ± S.E.M. *P < 0.05, **P < 0.01, ***P < 0.001, ****P < 0.001.

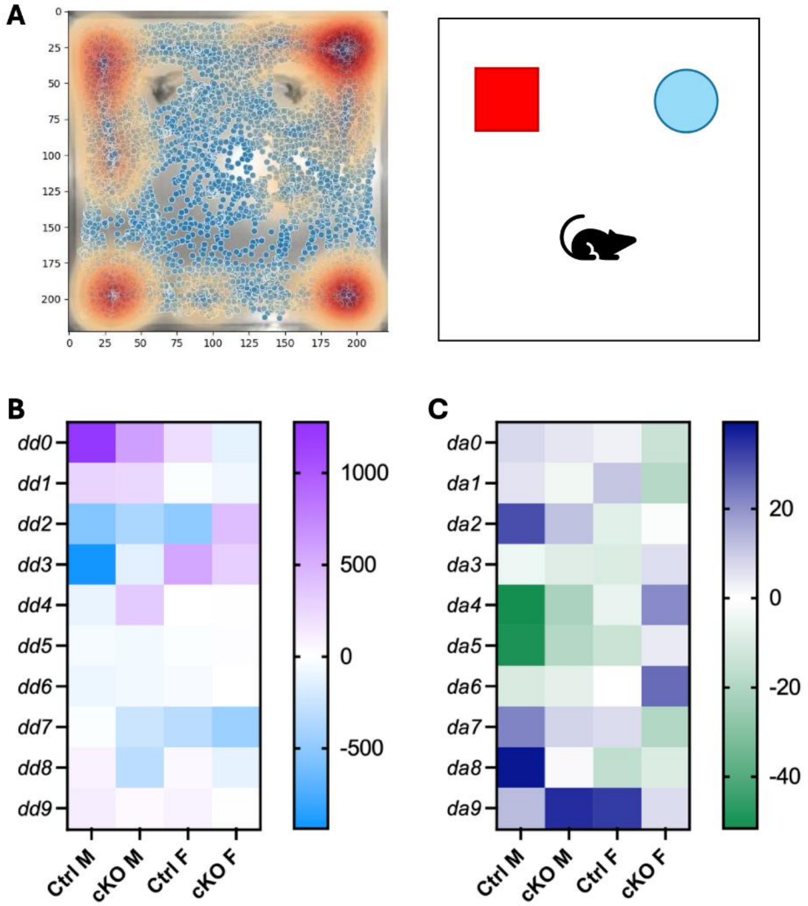


**Fig. S12. GR deletion in OPCs does not affect patterns of object exploration during novel object recognition test but reveal a general sex difference in the mouse positioning relative to the objects over time.** (**A**) Representative heatmap of arena and object exploration during novel object recognition test (NORT) and representation of NORT setup. (**B**) Heatmap comparing the normalized positioning distance of mice to objects during the testing phase of NORT (Three-way ANOVA: time $F(2.517, 146.0) = 3.113$, $p = 0.0362$; time x sex $F(9,522)=4.064$, $p<0.0001$; genotype $p>0.9999$). (**c**) Heatmap comparing the normalized turning angle of the mice during the testing phase of NORT (Three-way ANOVA: time x sex $F(9,522)=2.269$, $p=0.0169$; genotype $p>0.9999$).

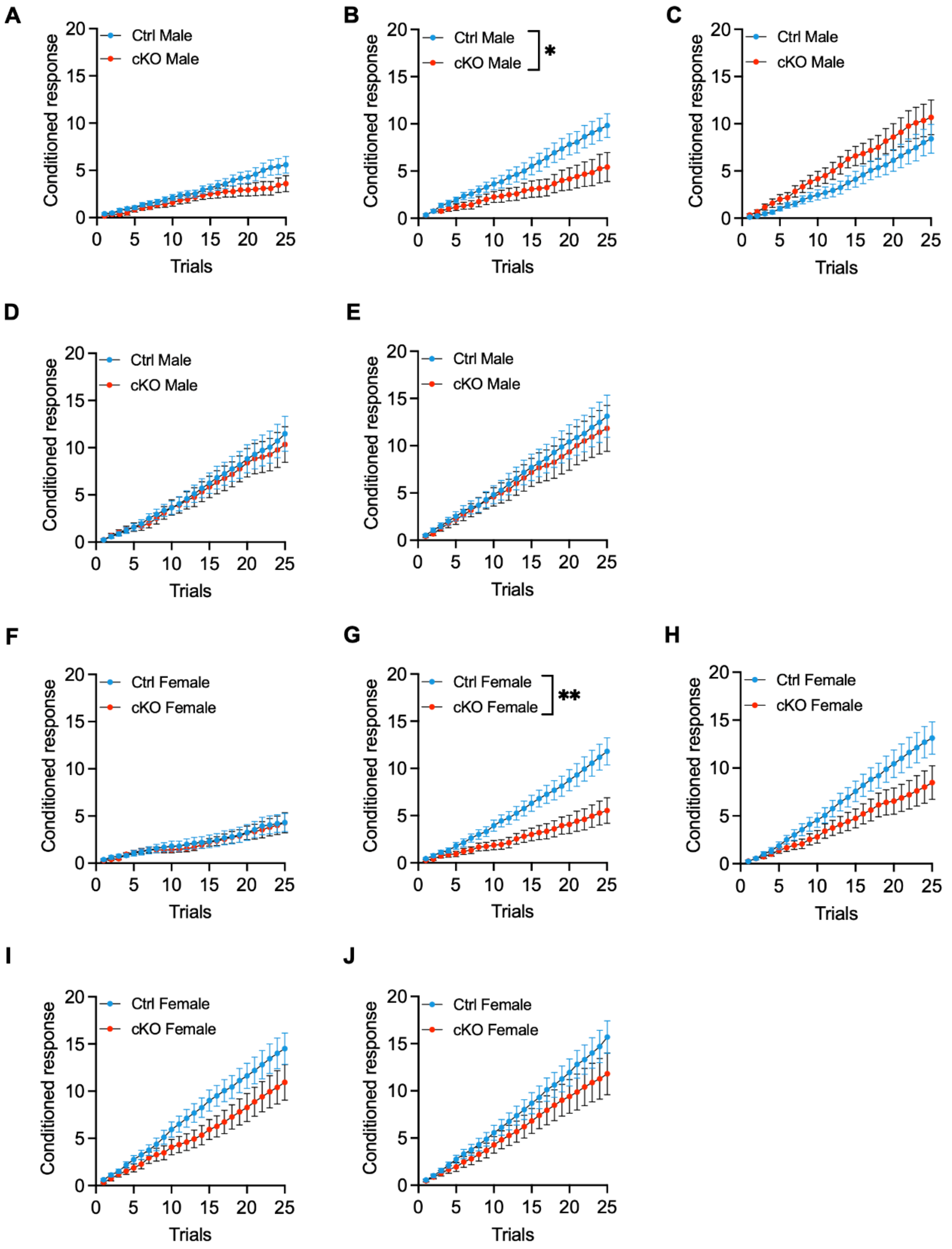
A
B
C
D
E
F
G
H
I
J
Conditioned response
Trials
Ctrl Male
cKO Male
Ctrl Female
cKO Female
*
**
0
5
10
15
20
25

**Fig. S13. GR deletion in OPCs affects aversive learning in both adult males and females.** Daily conditioned responses (CRs) across 25 trails over five consecutive days in the TWA test. (**A-E**) **Male mice**. (**A**) CR at day 1 (Two-way RM ANOVA: trial x genotype $F(1.351, 36.47) = 35.34$, $p = 0.0027$). (**B**) CR at day 2 (Two-way RM ANOVA: trial x genotype $F(24, 648) = 4.590$, $p < 0.0001$). (**C**) CR at day 3 (Two-way RM ANOVA, trial x genotype, $F(24, 648) = 0.4884$, $p < 0.0001$). (**D**) CR at day 4 (Two-way RM ANOVA: trial $F(1.108, 29.92) = 60.78$, $p < 0.0001$; genotype and interaction $p \varepsilon 0.79279$). (**E**) CR at day 5 (Two-way RM ANOVA: trial $F(24, 648) = 54.07$, $p < 0.0001$; genotype and interaction, $p \varepsilon 0.7426$). **(F-J) Female mice**. (**F**) CR at day 1 (Two-way RM ANOVA: trial $F(24, 696) = 21.03$, $p < 0.0001$; genotype and interaction, $p \varepsilon 0.06781$). (**G**) CR at day 2 (Two-way RM ANOVA: trial x genotype $F(24, 696) = 9.818$, $p < 0.0001$). (**H**) CR at day 3 (Two-way RM ANOVA: trial x genotype $F(24, 696) = 3.985$, $p < 0.0001$) . (**I**) CR at day 4 (Two-way RM ANOVA: trial x genotype $F(24, 696) = 2.516$, $p < 0.0001$). (**J**) CR at day 5 (Two-way RM ANOVA: trial $F(1.096, 31.79) = 90.33$, $p < 0.0001$; genotype and interaction, $p \varepsilon 0.0727$). Data are expressed as the mean ± S.E.M, *$P<0.05$, **$P < 0.01$, n = 17M/16F Ctrl and n = 12M/15F cKO.

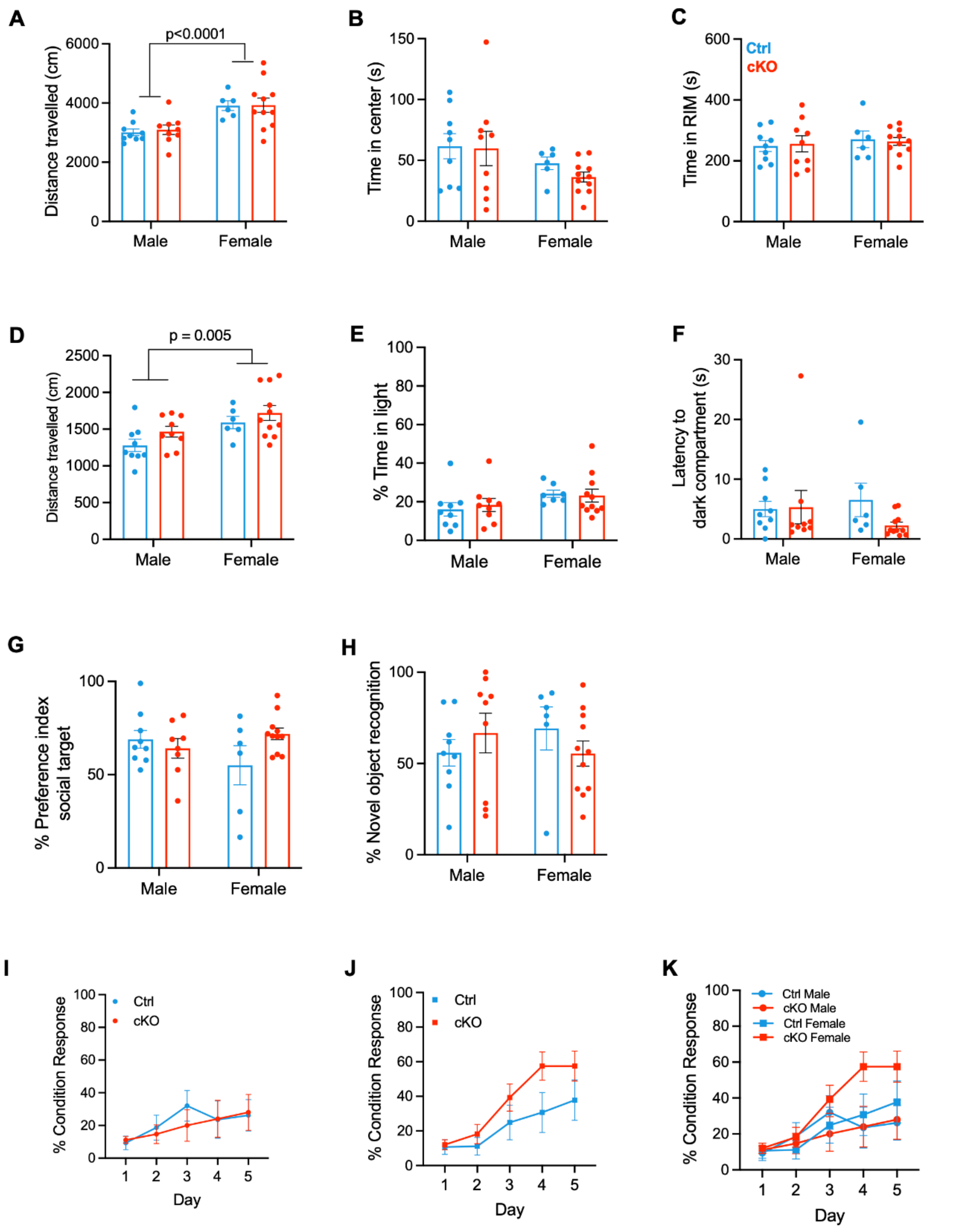


**Fig. S14. Deletion of GR in OPC in adulthood does not affect locomotion, cognitive performances, and anxiety-relevant behaviors.** (**A-C**) Open field test (OFT). (**A**) Distance travelled (two-way ANOVA, genotype, F (1, 31) = 0.07048, p=0.7924; sex, F (1, 31) = 19.53, p=0.00001; genotype x sex, F (1, 31) = 0.03289, p=0.8573). (**B**) Time in the center (two-way ANOVA, genotype, F (1, 31) = 0.4649, p=0.5004; sex, F (1, 31) = 3.687, p=0.0641; genotype x sex, F (1, 31) = 0.2312, p=0.6340). (**C**) Time in RIM (two-way ANOVA, genotype, F (1, 31) = 4.474e-007, p=0.9995; sex, F (1, 31) = 0.5076, p=0.4815; genotype x sex, F (1, 31) = 0.1065, p=0.7464). (**D-F**) Light-dark Box Test (LDBT). (**D**) Distance travelled (two-way ANOVA, genotype, F (1, 31) = 2.890, p=0.0991; sex, F (1, 31) = 9.151, p=0.005; genotype x sex, F (1, 31) = 0.09117, p=0.7647). (**E**) Percentage of time in the light compartment (two-way ANOVA, genotype, F (1, 32) = 0.04077, p=0.8413; sex, F (1, 32) = 3.895, p=0.0571; genotype x sex, F (1, 32) = 0.2490, p=0.6212). (**F**) Latency to the dark compartment (two-way ANOVA, genotype, F (1, 31) = 1.065, p=0.31; sex, F (1, 31) = 0.16, p=0.6919; genotype x sex, F (1, 31) = 1.438, p=0.2395). (**G**) Sociability/ Social interaction (two-way ANOVA, genotype, F (1, 30) = 1.158, p=0.2905; sex, F (1, 30) = 0.3074, p=0.5834; genotype x sex, F (1, 30) = 3.773, p=0.0615). (**H**) Percentage of time spent with a novel object (two-way ANOVA, genotype, F (1, 31) = 0.02386, p=0.8782; sex, F (1, 31) = 0.01255, p=0.9115; genotype x sex, F (1, 31) = 1.807, p=0.1886). (**I-K**) Two-way active avoidance. (**I**) Percentage of conditioned response over 5 days in Ctrl vs. cKO males (two-way RM ANOVA, genotype, F (1, 16) = 0.05503, p=0.8175; time, F (2.005, 32.08) = 3.425, p=0.0447; genotype x time, F (2.005, 32.08) = 0.5761, p=0.5682). (**J**) Percentage of conditioned response over 5 days in Ctrl vs. cKO females (two-way RM ANOVA, genotype, F (1, 18) = 2.330, p=0.1443; time, F (1.980, 35.63) = 19.79, p<0.0001; genotype x time, F (1.980, 35.63) = 1.824, p=0.1765). (**K**) Percentage of conditioned response over 5 days in Ctrl vs. cKO in males vs. females (three-way RM ANOVA, genotype, $F(1, 34) = 0.7066$, $p=0.4065$; sex, $F(1, 34) = 1.821$, $p=0.1861$; time, $F(2.101, 71.44) = 18.65$, $p<0.0001$; time x sex, $F(2.101, 71.44) = 3.918$, $p=0.0226$; time x genotype, $F(2.101, 71.44) = 1.238$, $p=0.2971$; sex x genotype, $F(1, 34) = 1.423$, $p=0.2411$; time x sex x genotype, $F(2.101, 71.44) = 1.112$; $p=0.3367$). Data are expressed as the mean ± S.E.M, *P<0.05, **P < 0.01, n = 9M/6F Ctrl and n = 9M/11F cKO.

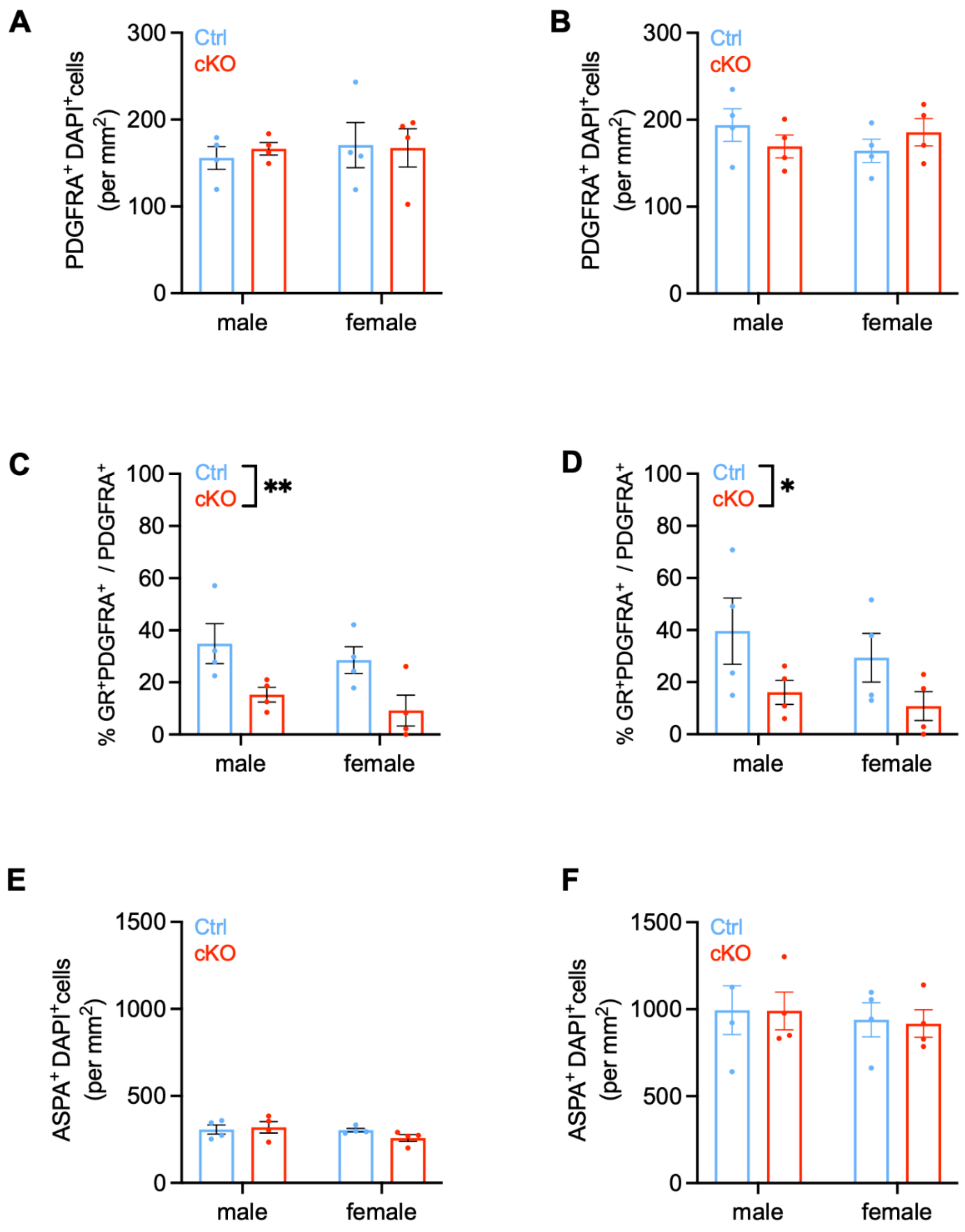


**Fig. S15. Deletion of GR in OPC in adulthood is comparably efficient in both sexes.** (**A**) OPC density in hippocampus (Two-way ANOVA, genotype, F (1, 12) = 0.03999, p=0.8448; sex, F (1, 12) = 0.1837, p=0.6758; genotype x sex, F (1, 12) = 0.1381, p=0.7167). (**B**) OPC density in cortex (Two-way ANOVA, genotype, F (1,

12) = 0.01109, p=0.9179; sex, F (1, 12) = 0.1854, p=0.6744; genotype x sex, F (1, 12) = 2.239, p=0.1604). (**C**) Percentage of GR+ OPCs in CA1 of the hippocampus (Two-way ANOVA, genotype, F (1, 12) = 11.89, p=0.0048; sex, F (1, 12) = 1.195, p=0.2958; genotype x sex, F (1, 12) = 0.0009083, p=0.9765). (**D**) Percentage of GR+ OPCs in the cortex (Two-way ANOVA, genotype, F (1, 12) = 5.891, p=0.0319; sex, F (1, 12) = 0.8020, p=0.3881; genotype x sex, F (1, 12) = 0.08102, p=0.7808. (**E**) OL density in hippocampus (Two-way ANOVA, genotype, F (1, 12) = 0.4844, p=0.4997; sex, F (1, 12) = 1.877, p=0.1958; genotype x sex, F (1, 12) = 1.410, p=0.2580). (**F**) OL density in cortex (Two-way ANOVA, genotype, F (1, 12) = 0.01408, p=0.9075; sex, F (1, 12) = 0.3486, p=0.5658; genotype x sex, F (1, 12) = 0.006014, p=0.9395).

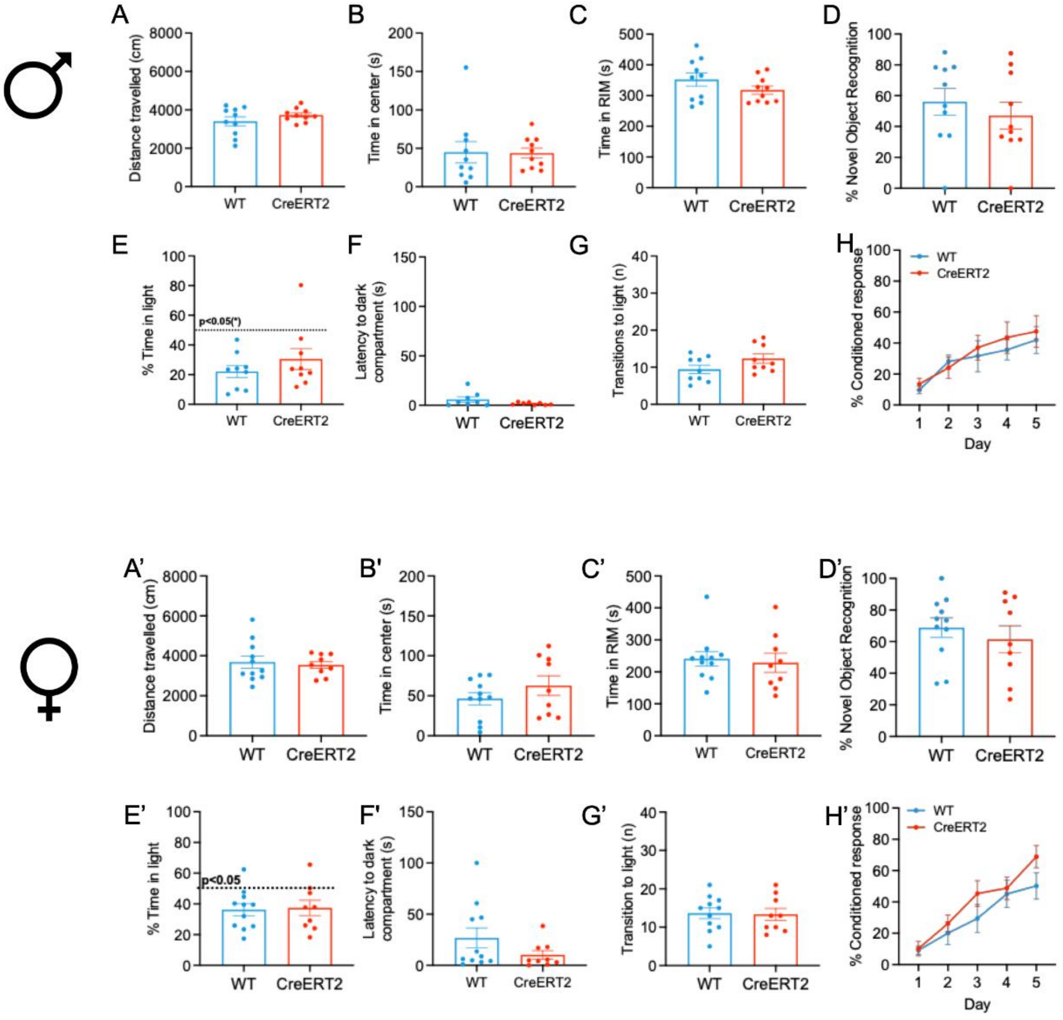


**Fig. S16. Lack of on NG2 allele does not alter behavioral readouts, which are instead affected by the loss of GR in NG2-glia in early life.** [GP1] (**A-H**) **Behavioral readouts in male mice**. (**A-C**) *Open field test:* (**A**) Locomotion (unpaired t test with Welch's correction: t = 1.247, df = 12.73, p = 0.2347). (**B**) Time spent in the centre of the arena (Mann-Whitney test: U=40.50, p = 0.4935). (**C**) Time spent in the rim of the arena (unpaired t test: t = 1.359, df = 18, p = 0.1909). (**D**) *Novel Object recognition test*: Percentage of time spent in the exploration of the new object normalised to the overall exploration time (unpaired t test: t = 0.7386, df = 18, p = 0.4696). (**E-G**) *Light-and-dark box test*: (**E**) Percentage of time spent in the illuminated compartment (Mann-Whitney test: U=33, p = 0.5457). (**F**) Latency to access the dark compartment (Mann-Whitney test: U=23, p = 0.3667). (**G**) Number of transitions to the illuminated compartment (unpaired t test: t = 1.732, df = 16, p = 0.1024). (**H**) *Two-way active avoidance*: Percentage of conditioned response over 5 consecutive days (Two-Way RM ANOVA: Day F (2.827, 45.23) = 14.48, p < 0.0001; genotype F (1, 16) = 0.1835, p 0.6741; day x genotype F (2.827, 45.23) = 0.4450, p = 0.7107). (**A'-H'**) **Behavioral readouts in female mice**. (**A'-C'**) *Open field test:* (**A'**) Locomotion (Unpaired t test: t = 0.3743, df=18, p = 0.7126). (**B'**) Time spent in the centre of the arena (Unpaired t test: t = 1.166, df=18, p = 0.2589). (**C'**) Time spent in the rim of the arena (Mann-Whitney test: U = 40, p = 0.5027). (**D'**) Percentage of time spent in the exploration of the new object normalised to the overall exploration time (Unpaired t test: t = 0.7117, df=18, p = 0.4858). (**E'-G'**) *Light-and-dark box test*: (**E'**) Percentage of time spent in the illuminated compartment (Unpaired t test: t = 0.1793, df=18, p = 0.8597). (**F'**) Latency to access the dark compartment (Mann-Whitney test: U =32, p = 0.2014. (**G'**) Number of transitions to the illuminated compartment (Unpaired t test: t = 0.1459, df=18, p = 0.8856). (**H'**) *Two-way active avoidance*: Percentage of conditioned response over 5 consecutive days (Two-Way RM ANOVA: Day F (2.828, 50.90) = 34.70, p < 0.0001; genotype F (1, 18) = 1.171, p 0.2934; day x genotype F (2.828, 50.90) = 1.354, p = 0.2678). Data are expressed as the mean

± S.E.M, *P<0.05, **P < 0.01, *Males:* n = 8-10 WT and n = 8-10 CreERT2; *Females*: n = 11 WT and n = 9 CreERT2.
